# Development of a bioassay guided genome mining approach for antifungal natural product discovery from pseudomonads

**DOI:** 10.1101/2025.05.30.657047

**Authors:** George Lund, Susan Mosquito, David M Withall, John Caulfield, David Hughes, Ian M Clark, Jason Rudd, Tim H Mauchline

## Abstract

*Zymoseptoria tritici* causes Septoria Leaf Blotch disease of wheat and has evolved to overcome most chemical and genetic control methods. As such, new tools are required for future disease control.

We identified *Pseudomonas* isolates that antagonise *Z. tritici* through the production of secreted secondary metabolites, using a novel *in vitro Z. tritici* antagonism assay. In addition to high-throughput qualitative assessment of *Pseudomonas* antagonism of *Z. tritici*, a quantitative assessment identified variation in the sensitivity of *Z. tritici* isolates to antagonism by a subset of *Pseudomonas* isolates.

Genome assemblies of 3 strongly antagonistic *Pseudomonas* isolates were found to contain a predicted Biosynthetic Gene Cluster (BGC) with high sequence similarity to a reference BGC encoding the biosynthesis of the known antifungal compound 2,4-diacetylphloroglucinol (2,4-DAPG).

Mutagenesis of the core biosynthetic gene *phlD* resulted in a loss of 2,4-DAPG production in *Pseudomonas* isolate Roth82, and a loss of *Z. tritici* inhibition in the antagonism assay. These results demonstrate that the described *in vitro* antagonism assay can be used to identify, quantify and mechanistically characterise bacterial antagonism of *Z. tritici* through the production of secondary metabolites.

This is the first study to find significant differences in the response of genetically diverse isolates of *Z. tritici* isolates to bacterial antagonists, suggesting sensitivity to bacterial antagonism exists as a quantitative trait within natural *Z. tritici* populations. Our approach can be used to identify and characterise putatively novel BGCs that encode natural products with antifungal activity against *Z. tritici*.

## Introduction

Wheat is the UK’s most widely grown crop, but yield is diminished due to diseases caused by pathogenic fungi. The most important disease of wheat in the UK is Septoria tritici blotch, caused by the fungus *Zymoseptoria tritici*, resulting in up to 50% yield losses (Torriani, 2015). Wheat has no durable broad ranging natural resistance to infection and the fungus has become insensitive to all known classes of existing commercial fungicides (Lucas et al., 2015). Therefore, new ways to protect wheat are urgently needed. One underexplored approach is to exploit the natural antifungal chemistries that are encoded in microbial genomes.

The soil microbiome is hugely diverse and largely untapped for the discovery of new molecules that can suppress plant pathogens. To date, most antimicrobial compounds have been derived from soilborne actinomycetes (Barka et al., 2016). *Pseudomonas* bacteria are an important component of the soil microbiome with exemplar species behaving as human pathogens(Kerr and Snelling, 2009) (e.g. *P. aeruginosa*), plant pathogens (Morris et al., 2013) (e.g. *P. syringae*), as well as plant growth promoting bacteria (Santoyo, 2016), and biocontrol agents (Höfte, 2021; Thomashow and Weller, 1996) (e.g. *P. fluorescens*); yet the diverse secondary metabolite repertoire of this genus, which has potential for the discovery of new high value fungicides, is relatively underexplored experimentally (Saati-Santamaría et al., 2022).

Previous studies investigating *in vitro* antagonism of *Z. tritici* by *Pseudomonas aeruginosa* isolate LEC1, found the siderophore pyocyanin (Flaishman et al., 1990) and 1-hydroxyphenazine (Levy et al., 1989, 1992a) to partially contribute towards antagonism. *Pseudomonas fluorescens* PMF2 was found to antagonise *Z. tritici* through the production of 2,4-diacetylphloroglucinol (2,4-DAPG) and 2 additional unidentified antifungal compounds (Levy et al., 1992b). Tropolone, produced by *Pseudomonas lindbergii*, was found to inhibit the growth of *Z. tritici in vitro* (Levy et al., 1988; Lindberg et al., 1980). *Pseudomonas* isolates have also previously been shown to antagonise *Z. tritici* through the production of volatile compounds, with *Pseudomonas putida* BK8661 found to antagonise *Z. tritici* through the production of hydrogen cyanide (HCN) *in vitro* (Flaishman, 1996). *In vitro* studies investigating the antagonistic potential of bacterial isolates against fungal plant pathogens, have mostly consisted of single genotypes of fungal plant pathogens (Flaishman et al., 1990; Flaishman, 1996; Levy et al., 1992a, 1992b, 1989, 1988). However, *Z. tritici* has high levels of genetic diversity, with previous studies identifying over 600 different genotypes of the fungus in a single field over a single season (Chen and McDonald, 1996), and individual leaf lesions were found to harbour 2-5 different genotypes (Linde et al., 2002).

Quantification of *in vitro* antagonism of fungal pathogens by bacterial isolates is difficult, owing to the filamentous hyphal growth of many fungal pathogens, and therefore measurements have commonly taken place through qualitative assessments (Bubici et al., 2019; Mauchline et al., 2015). Quantitative assessment of *in vitro Z. tritici* antagonism could facilitate the identification of *Pseudomonas* isolates capable of antagonising *Z. tritici in vitro* in a broader context. Such assessments could facilitate sensitivity comparisons of genetically diverse *Z. tritici* isolates to antagonistic *Pseudomonas* isolates. Pseudomonads are amenable to culture and this property facilitates the acquisition of their genomes, which in turn assists in the identification of the genetic components responsible for the biosynthesis of antifungal or fungistatic secondary metabolites (Gross and Loper, 2009).

This study introduces a novel qualitative and quantitative blastopore-based assay for high throughput assessment of pseudomonad growth suppression of multiple *Z. tritici* genotypes. The radial zones of clearing produced by antagonistic *Pseudomonas* isolates, confronted with *Z. tritici* blastospores as a confluent lawn, offer the potential to both qualitatively and quantitatively assess antagonism of *Z. tritici* by *Pseudomonas* isolates. This assay in combination with genomics, microbial mutagenesis and analytical chemistry enables the prioritisation of predicted secondary metabolite Biosynthetic Gene Clusters (BGCs) from antagonistic isolates for further characterisation, and comparison with BGCs previously described in literature from the MIBiG 3.0 repository (Terlouw et al., 2023). We report the initial screening of a *Pseudomonas* library of 534 isolates for suppression of the *Z. tritici* reference isolate IPO323. Subsequent *in vitro* testing of 5 antagonistic and 6 non-antagonistic *Pseudomonas* isolates confronted with 12 genetically diverse *Z. tritici* isolates, revealed significant differences in resistance to bacterial antagonism. To our knowledge this is the first study to find evidence suggesting sensitivity to bacterial antagonism exists as a quantitative trait within natural *Z. tritici* populations. We demonstrate the power of a bioassay-mediated genomics approach to define potential antifungal secondary metabolite BGCs involved in bacterial-fungal interactions *in vitro*. The study also demonstrates the importance of testing genetically diverse individuals from within pathogen populations to ensure consistency of antagonistic responses ahead of further exploitation of bacterial secondary metabolites or biological control agents as future disease control strategies.

## Results

### Qualitative screening of *Z. tritici* IPO323 antagonism

A total of 534 *Pseudomonas* isolates were screened qualitatively for antagonism of *Z. tritici* IPO323 blastospores, with 52 identified as being antagonistic (Figure 1, Table S1).

**Figure 1.**
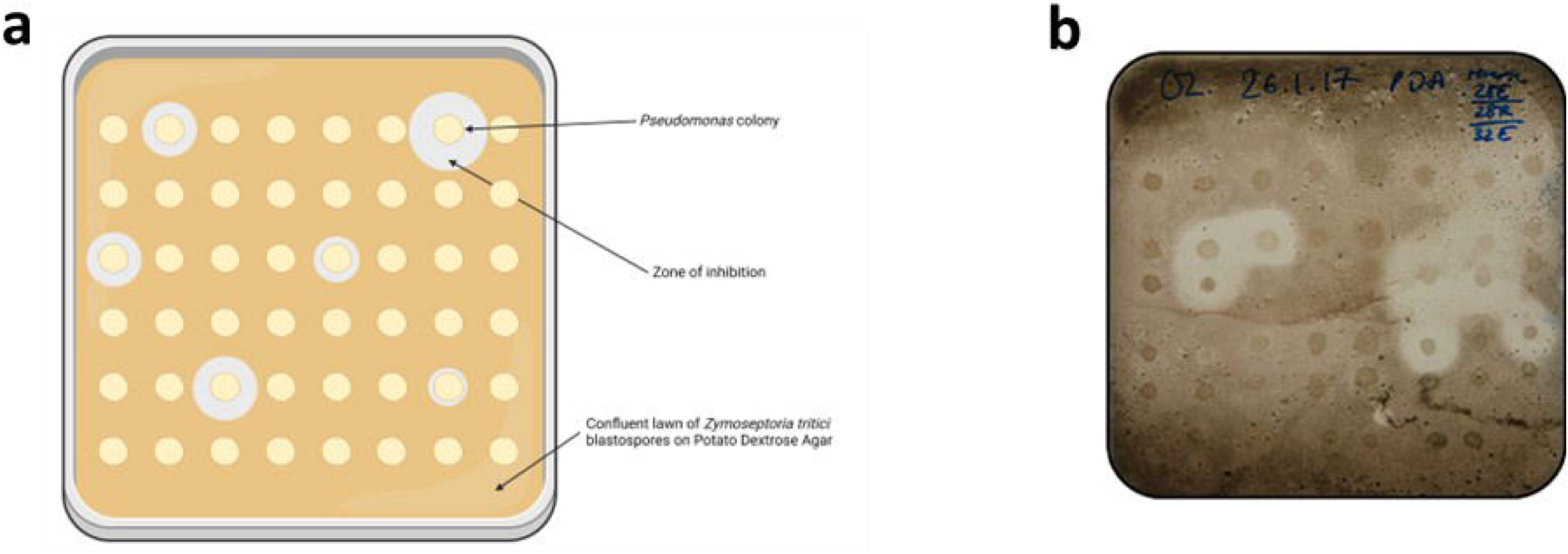
A high-throughput qualitative assay to assess the ability of Pseudomonas antagonism of Z. tritici blastospores in vitro. **(a)** Schematic diagram of the qualitative *Z. tritici* blastopore antagonism assay. In the bioassay, *Pseudomonas* isolates are spotted onto a confluent lawn of *Z. tritici* blastospores on PDA. Created in BioRender. Lund, G. (2024) BioRender.com/p07z107 **(b)** A photograph of the qualitative *Z. tritici* blastospore antagonism assay, with 48 *Pseudomonas* colonies on one assay plate in confrontation with a single fungal isolate.

*Pseudomonas* isolates demonstrate variable quantitative suppression of a genetically diverse *Z. tritici* collection.

Of the 52 *Pseudomonas* isolates that demonstrated qualitative antagonism to *Z. tritici* IPO323 blastospores, 5 antagonistic isolates alongside 6 non-antagonistic isolates (Table S2) from the collection were randomly selected and screened quantitatively for antagonism against a collection of 12 genetically diverse *Z. tritici* isolates isolated from across Europe in 2015/16 (Chen et al., 2023) (Figure 2). All 5 qualitatively antagonistic *Pseudomonas* isolates were found to suppress growth of *Z. tritici* IPO323 as well as the additional 11 genetically diverse *Z. tritici* isolates in the quantitative antagonism assay (Figure 2). In addition, all 6 *Pseudomonas* isolates which had been determined to be qualitatively non-antagonistic towards *Z. tritici* IPO323, as well as LB Miller Broth medium controls, did not produce an observable zone of clearing when confronted with any of the *Z. tritici* isolates screened. This indicates that IPO323 confrontation is an appropriate starting test for the identification of *Pseudomonas* isolates with the potential to antagonise *Z. tritici* isolates qualitatively *in vitro*.

**Figure 2.**
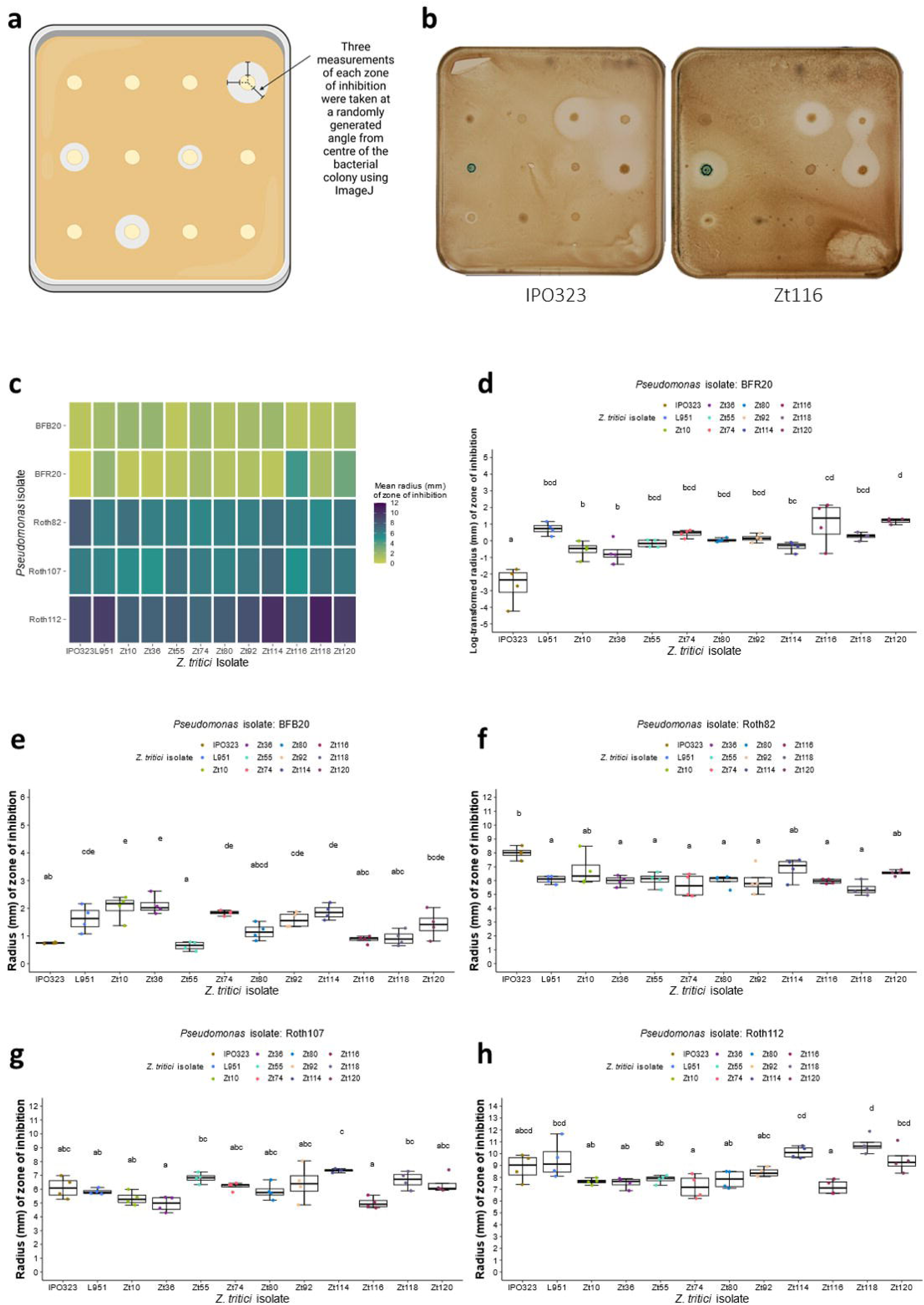
Significant differences in zones of inhibition of genetically distinct*Z. tritici* blastospores by *Pseudomonas* isolates *in vitro*, using a quantitative antagonism assay. (**a**) A schematic diagram of the quantitative *Z. tritici* blastopore antagonism assay. Created in BioRender. Lund, G. (2023) BioRender.com/y78z760 **(b)** Images of *in vitro* quantitative antagonism plates exhibiting differences in zones of inhibition produced by *Pseudomonas* isolates against IPO323 and Zt116 blastospores. **(c)** Heatmap indicating the average radius of the zone of inhibition produced in each pairwise interaction between antagonistic *Pseudomonas* and *Z. tritici* isolates in the quantitative antagonism assay after 12 days (n=4). **(d-h)** Zones of inhibition produced by *Pseudomonas* isolates in confrontation with genetically diverse *Z. tritici* blastospores *in vitro*. Mean values of zones of inhibition not sharing any letter are significantly different by Sidak’s test for multiple comparisons at the 5% level of significance for *Pseudomonas* isolates **(d)** BFR20 **(e)** BFB20 **(f)** Roth82 **(g)** Roth107 **(h)** Roth112.

A statistically significant difference in the mean zones of inhibition (millimetres) was detected between genetically diverse *Z. tritici* isolates confronted with antagonistic *Pseudomonas* isolates; BFR20 (*F*(11, 36) = 11.48, *p* < 0.001), BFB20 (*F*(11, 36) = 10.97, *p* < 0.001), Roth82 (*F*(11, 36) = 4.455, *p* < 0.001), Roth107 (*F*(11, 36) = 5.567, *p* < 0.001), and Roth112 (*F*(11, 36) = 8.11, *p* < 0.001). Significant differences in the mean zones of inhibition produced by antagonistic *Pseudomonas* were determined using Sidak’s multiple comparison post-hoc test (Figure 2 d-h, Tables S4-S13).

### Genome sequencing, taxonomic estimation, and predictive bioinformatics of secondary metabolism of environmental *Pseudomonas* isolates

Genome assemblies were generated by MicrobesNG for the 11 *Pseudomonas* isolates examined in the quantitative *Z. tritici* antagonism assay. All generated genome assemblies were found to have BUSCO scores >99%, when analysed using the bacterial odb10 dataset (containing 124 single copy orthologous genes) to check for bacterial contamination within the genome assembly (Figure S1). Additionally, BUSCO analysis of the generated genome assemblies using the Pseudomonadales odb10 dataset (containing 782 single copy orthologous genes) also returned BUSCO scores >99% for all assemblies. As such both BUSCO analyses indicated high levels of genome assembly completeness and low levels of genome assembly contamination (Figure S2).

To predict putative secondary metabolite biosynthetic gene clusters from the *Pseudomonas* genome assemblies, antiSMASH 7.0.0 (Blin et al., 2023) was used. A total of 131 BGCs were predicted from the genome assemblies of the 11 *Pseudomonas* isolates (Figure S3). To compare predicted BGCs between *Pseudomonas* isolates, and compare to reference BGCs from the MIBiG 3.0 database (Terlouw et al., 2023), sequence similarity networks of predicted secondary metabolite BGCs from both antagonistic and non-antagonistic *Pseudomonas* isolates were generated using BiG-SCAPE (Navarro-Muñoz et al., 2020), including reference BGCs from MIBiG 3.0. A total of 46 GCF networks were formed of two or more BGCs, with 3 GCFs containing a reference BGC from MIBiG 3.0 (GCF_00413, GCF_02527, GCF_02551, Figure 3a), indicating similarity of predicted BGCs from *Pseudomonas* genome assemblies within this study, at the 0.3 threshold in BiG-SCAPE. A total of 13 GCF networks were found to contain predicted BGCs from solely *Z. tritici* blastospore antagonistic *Pseudomonas* isolates (Figure 3a). Of these, GCF_02551 contained reference BGCs from MIBiG 3.0, BGC0000280 and BGC000281, which have been shown experimentally to encode the biosynthesis of 2,4-diacetylphloroglucinol (Bangera and Thomashow, 1999), a known antifungal compound effective at inhibiting *Z. tritici in vitro* (Levy et al., 1992b). Predicted BGCs within GCF_02551 were aligned using clinker, with high levels of amino acid similarity, and synteny identified between the reference BGCs and those predicted from the generated *Pseudomonas* genome assemblies (Figure 3b).

**Figure 3.**
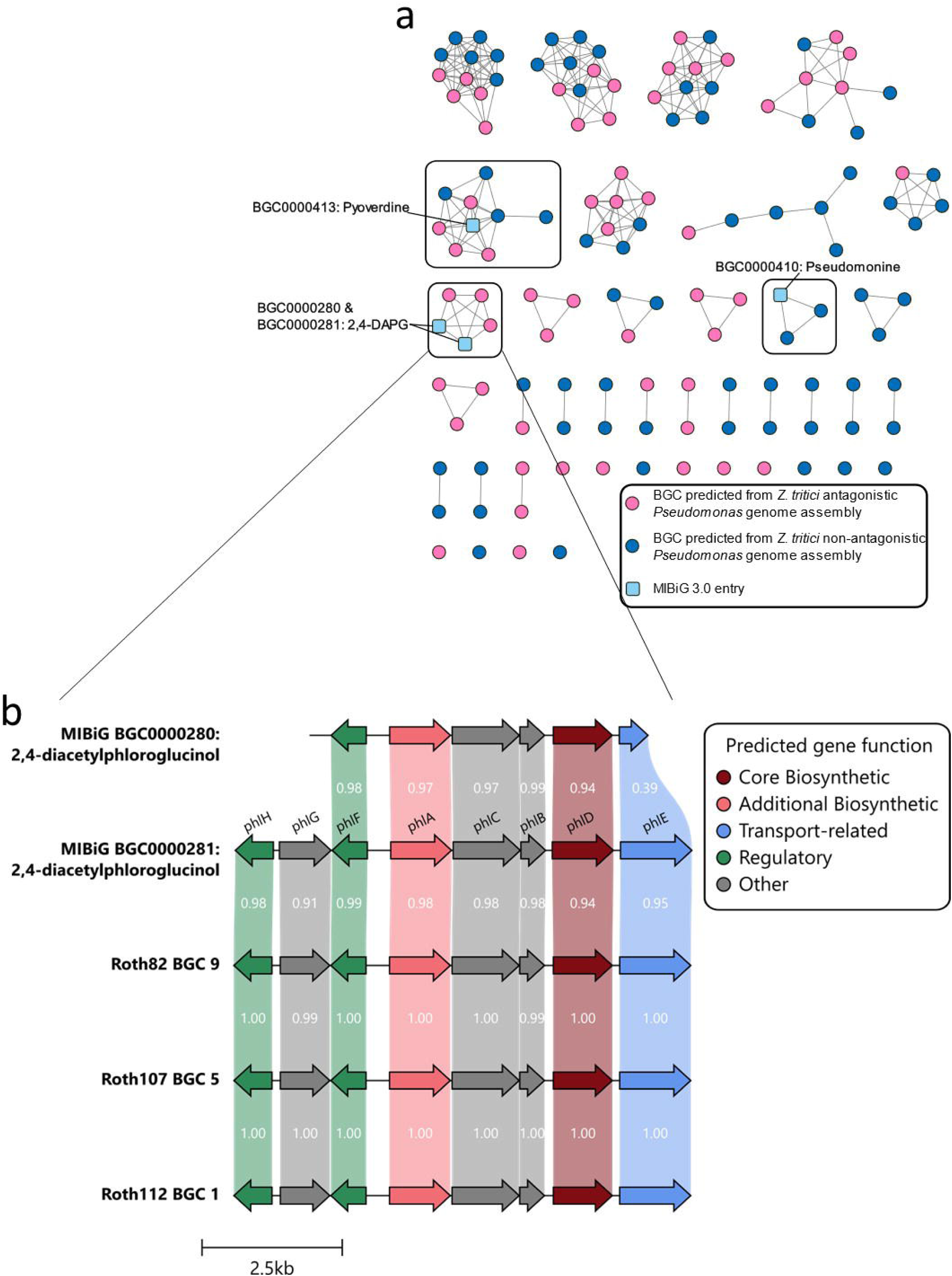
Sequence similarity networks generated using BiG-SCAPE reveal Gene Cluster Families associated with the *Z. tritici* blastospore antagonism phenotype, including reference BGCs with known antifungal activity from the MIBiG repository. (**a**) Sequence similarity networks indicated predicted BGCs with high sequence similarity to MIBiG reference BGCs for pyoverdine (BGC0000413, GCF_02527), 2,4-DAPG (BGC0000280 & BGC0000281, GCF_02551), and pseudomonine (BGC0000410, GCF_00413), within the *Pseudomonas* genome assemblies. (**b**) Amino acid alignment of BGCs forming GCF_02551 with 2,4-DAPG biosynthesis reference BGCs from MIBiG (GC000280 & BGC000281) indicates high levels of amino acid similarity and gene synteny between predicted BGCs.

To estimate the taxonomy of the generated *Pseudomonas* genome assemblies, autoMLST (Alanjary et al., 2019) was used. A phylogenetic tree was constructed using 100 common genes (Table S3) from the 11 *Pseudomonas* genome assemblies, with the closest 50 genome assemblies from RefSeq included in the phylogenetic tree construction. The closest RefSeq type strains to the genome assemblies generated in this study were cross referenced with previous literature to identify clades associated with described subgroups of the *P. fluorescens* species complex (Hesse et al., 2018; Lauritsen et al., 2021) (Figure 4).

**Figure 4.**
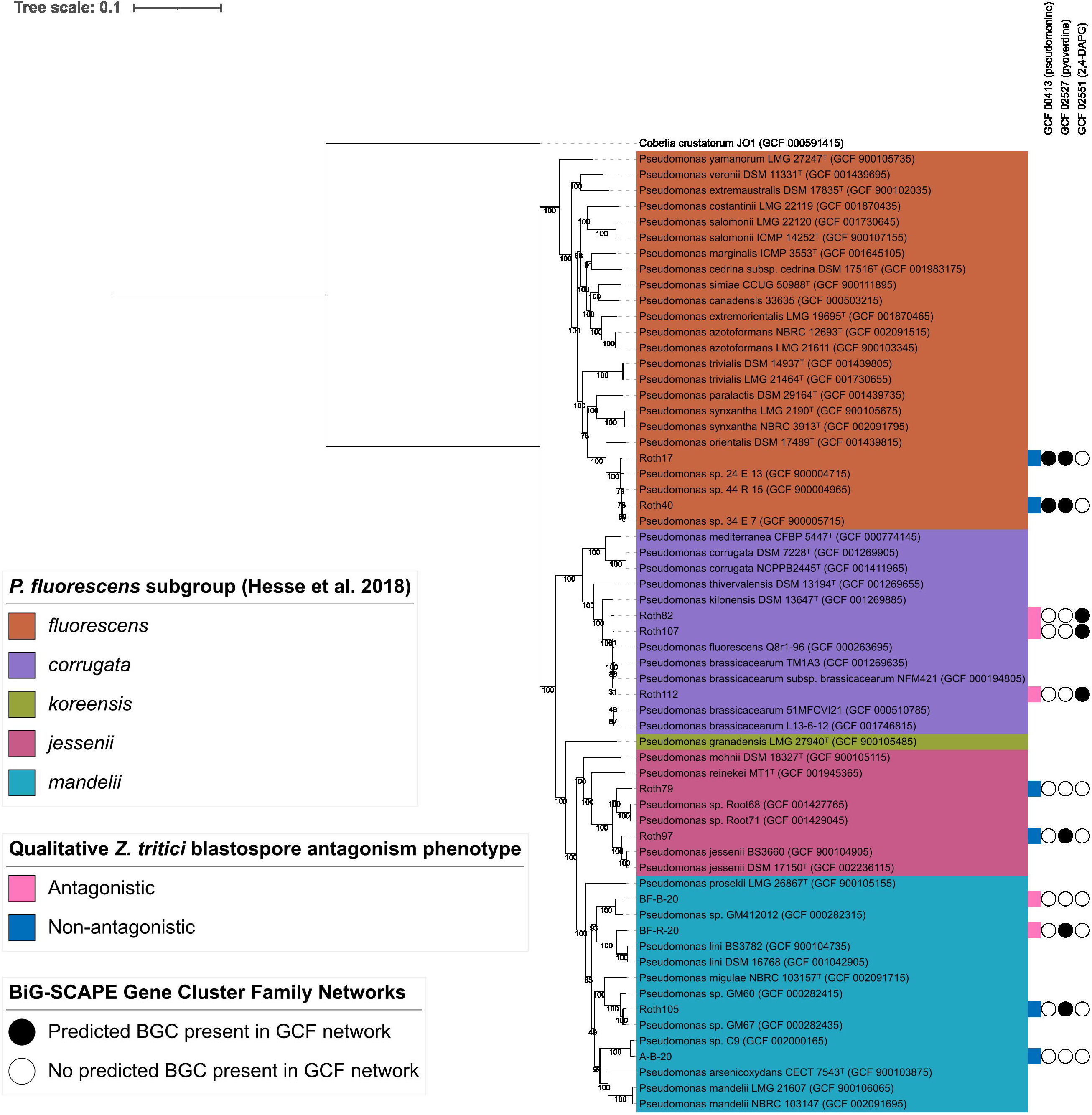
Phylogenetic distribution of *Pseudomonas* isolates with predicted secondary metabolite BGCs with significant similarity to known BGCs in relation to*Z. tritici* blastopore antagonism phenotypes. Phylogenetic analysis with known *Pseudomonas* strains, using autoMLST, putatively identifies 11 *Pseudomonas* strains used within this study within the *P. fluorescens*, *P. corrugata*, *P. koreensis*, *P. jessenii*, and *P. mandelii* subgroups of the *P. fluorescens* species complex. The presence and absence of predicted BGCs that form sequence similarity networks with experimentally characterised BGCs from the MIBiG database are indicated, alongside qualitative *Z. tritici* blastospore antagonism phenotype against *Z. tritici* strain IPO323. Type strain *Pseudomonas* genomes identified according to LPSN (accessed 24 January 2025) are denoted by a superscript ‘T’.

Phylogenetic analysis indicated that the 11 *Pseudomonas* strains within this study span 4 distinct subgroups of the *P. fluorescens* species complex, as defined by Hesse *et al* (Hesse et al., 2018). Isolates predicted within the *P. corrugata* subgroup, Roth82, Roth107 and Roth112, were all found to exhibit antagonistic phenotypes when confronted with *Z. tritici* blastopores *in vitro*. Both *Z. tritici* blastospore antagonistic (BF-R-20) and non-antagonistic (A-B-20, Roth105) *Pseudomonas* isolates were found within the *P. mandelii* subgroup. Isolates Roth17 and Roth40, within the *P. fluorescens* subgroup, and Roth79 and Roth97 within the *P. jessenii* subgroup were all found to exhibit a non-antagonistic phenotype when confronted with *Z. tritici* blastospores *in vitro*.

The resulting phylogenetic tree was annotated with the GCF network data generated using BiG-SCAPE, to visualise the distribution of predicted BGCs with a high similarity to MIBiG reference BGCs (Figure 4), and all predicted BGCs forming GCF networks (Figure S5). All 3 isolates classified within the *P. corrugata* subgroup (Roth82, Roth107 and Roth112) were found to contain predicted BGCs that formed a sequence similarity network with MIBiG reference BGCs encoding the production of 2,4-DAPG. The genome assemblies of *Z. tritici* non-antagonistic *P. fluorescens* subgroup isolates Roth17 and Roth40 were found to contain predicted BGCs with significant similarity to the MIBiG reference BGC encoding the biosynthesis of pseudomonine. Predicted BGCs with a significant similarity to MIBiG reference BGC BGC0000413, encoding the production of pyoverdine, were found in genome assemblies of the non-antagonistic *P. fluorescens* subgroup (Roth17 & Roth40), *P. jessenii* subgroup (Roth97), and *P. mandelii* subgroup (Roth105), as well as the antagonistic *P. mandelii* subgroup (BF-R-20) isolate.

### Gene disruption of BGC predicted to encode the biosynthesis of 2,4-DAPG results in a loss of visible *Zymoseptoria tritici* antagonism, and a reduction in 2,4-DAPG detected in agar extracts of confrontation zones

To examine the involvement of known antifungal secondary metabolite 2,4-DAPG in the observed *Z. tritici* blastospore antagonism by isolate Roth82 (*P. corrugata* subgroup), a Δ*phlD* mutant was constructed by allelic replacement, in which the *phlD* gene was disrupted in Roth82, resulting in Roth82Δ*phlD* (Figure S6). *Zymoseptoria tritici* blastospore confrontation assays were undertaken with strain IPO323, with methanol agar extracts from the zone surrounding Roth82 and Roth82Δ*phlD* prepared after 12 days of confrontation. No visible zone of inhibition was present for Roth82Δ*phlD* (Figure 5b), whereas a zone of inhibition was seen for wild-type Roth82 (Figure 5a). These observed phenotypes correlated with LC-MS analysis of the agar extracts, with wild-type Roth82 extract showing a peak at *m/z* 209.0455 ± 5ppm [M-H]^-^ with a retention time of 26.65 minutes (Figure 5c), corresponding to the authentic chemical standard for 2,4-DAPG (Sigma, Figure 5c). For the Roth82Δ*phlD* mutant confronted with *Z. tritici* blastospores, no peak corresponding to the retention time and *m/z* 209.0455 [M-H]^-^ of the authentic chemical standard of 2,4-DAPG was detected (Figure 5c).

**Figure 5.**
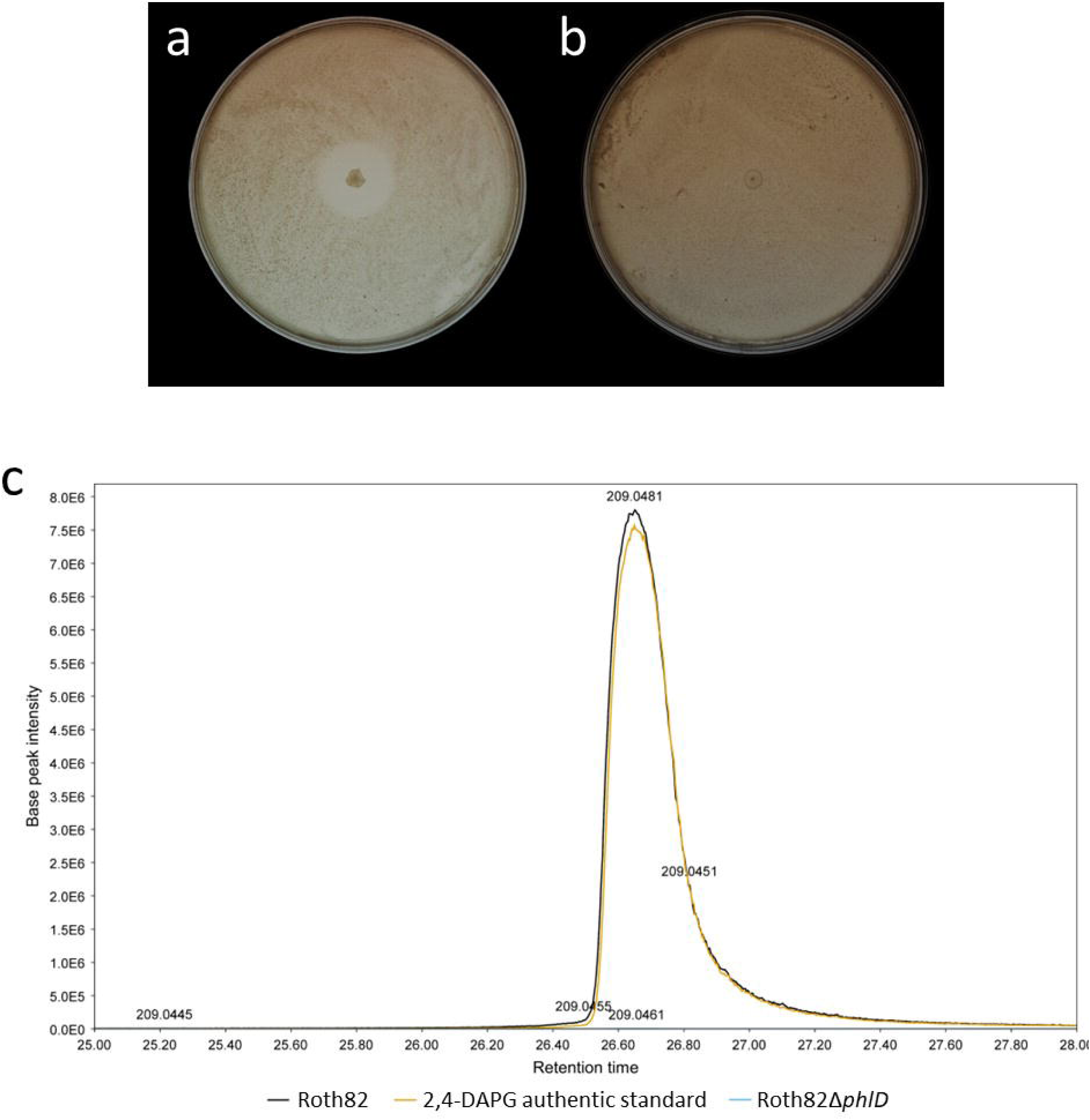
Disruption of *phlD* in isolate Roth82 (*P. corrugata* subgroup) results in a loss of *Z. tritici* blastospore antagonism *in vitro*, and reduction in the production of 2,4-DAPG. *Zymoseptoria tritici* IPO323 blastospore antagonism assay with wild-type Roth82 exhibiting a zone of inhibition (a) and Roth82Δ*phlD* showing no zone of inhibition (b). (c) Extracted ion chromatogram at m/z = 209.0455 with a mass tolerance of ± 5ppm, corresponding to [M-H] (molecular formula C_10_H_9_O_5_) of 2,4-DAPG (C_10_H_10_O_5_). Liquid Chromatography-Mass Spectrometry analysis was performed on agar extracts from the area surrounding the colony of wild-type Roth82 in confrontation with *Z. tritici* IPO323 blastospores (black), Roth82Δ*phlD* in confrontation with *Z. tritici* IPO323 blastospores (turquoise), and the authentic chemical standard for 2,4-DAPG (Sigma, orange).

## Discussion

Our approach utilised a novel high throughput phenotyping screening methodology to identify environmental *Pseudomonas* isolates with an ability to suppress growth of the wheat foliar pathogen *Z. tritici* from a culture collection. We combined *in vitro* phenotyping method with genome mining of antagonistic *Pseudomonas* isolates to identify putative antifungal secondary metabolite biosynthetic gene clusters and associated gene cluster families. This led to the identification of 52 isolates from a collection of 534 with an ability to suppress *Z. tritici* IPO323 blastospores *in vitro.* Genome sequencing and assembly of a subset of this collection, comprising 5 antagonistic and 6 non-antagonistic strains was conducted, followed by the prediction of 131 BGCs that formed 46 discrete GCF networks.

In addition to demonstrating high throughput qualitative screening, our results also indicate that the blastospore antagonism assay can be used to quantitatively assess antagonism of *Z. tritici* growth by *Pseudomonas* isolates *in vitro*, albeit with lower throughput due to increased spacing of bacterial cultures on test plates and the requirement to adopt digital image analysis to quantify inhibition zones. We believe that this assay can be expanded to assess the ability of other microbial species to suppress *Z. tritici* growth, though we recommend that this is not done in a disparate manner (e.g., wholesale screening of bacterial culture collections), but rather tailored toward specific bacterial genera to reduce quality control issues which would likely be encountered when simultaneously screening bacteria with inherently different growth kinetics.

Assessment of fungal antagonism by bacterial isolates is commonly conducted qualitatively, owing to the difficulty in quantifying antagonism of hyphal networks growing across entire plates *in vitro*. However, a high throughput semi-solid medium based screen has been described, using spectrophotometer measurements (OD_600_) and turbidimetric measurements to quantify growth of the pathogenic fungus *Aspergillus flavus* when exposed to chemical libraries, to screen for antagonism of the fungus through a reduction in growth rate compared to control (Medina et al., 2012). However, the application of this technique is limited for our purposes, due to the overlapping spectra commonly used to quantify the growth of bacteria and *Z. tritici*, hampering the discrimination of bacterial isolates and *Z. tritici* growth.

This is the first study to find significant differences in the response of genetically diverse isolates of a *Z. tritici* to bacterial antagonists. Our ability to identify subtle differences in the *in vitro* growth profiles of *Z. tritici* in a quantitative manner, when confronted with a given antagonistic pseudomonad, indicates that there is natural variation in sensitivity or resistance to bacterial antagonists and associated secondary metabolites among *Z. tritici* isolates obtained from across Europe. We found that significant differences in the variability of inhibition between the original *Z. tritici* reference isolate IPO323 (isolated in 1984) and the more recently isolated strains were observed in response to confrontation with some pseudomonads, such as BFR20. This suggests that selection for broad spectrum antifungal insensitivity in *Z. tritici* across this period, may be correlated with differences in response to in vitro bacterial antagonism and further studies to identify mechanistic understanding of this phenomenon are required.

A number of quantitative traits have been identified in *Z. tritici* populations, for traits such as temperature tolerance (Lendenmann et al., 2016), and fungicide resistance (Lendenmann et al., 2015). The results presented within this study highlight the possibility of sensitivity to bacterial antagonism may also exist as a quantitative trait within natural *Z. tritici* populations. The patterns of sensitivity of *Z. tritici* isolates to antagonism by *Pseudomonas* isolates show that no *Z. tritici* isolate tested was consistently the most sensitive or resistant towards all antagonistic *Pseudomonas* isolates screened. Our antagonism phenotype paired bioinformatic analyses suggest that this could indicate differing underlying mechanisms of antagonism by the *Pseudomonas* isolates, and potentially differing genetic targets for antagonism in *Z. tritici* blastospores, or differing levels of tolerance to antifungal metabolites in the *Z. tritici* test set. In future work it will be interesting to generate mapping populations of sexual crosses between *Z. tritici* isolates resistant and/or sensitive to particular bacterial antagonists, to identify possible ‘modes-of-action’ of bacterial antagonists. This could be achieved through the screening of the mapping population in the blastospore confrontation assay, and the use of Genome-Wide Association Study (GWAS) approaches to identify genomic regions in *Z. tritici* associated with heightened sensitivity or resistance to antagonism.

We acknowledge that a limitation of our approach employed in this study, is that the observed zones of clearing produced by antagonistic *Pseudomonas* isolates are potentially constrained by the diffusion of secreted compounds involved in antagonism through the agar medium. This could limit the ability of this screen to identify subtle changes in sensitivity of *Z. tritici* isolates to classes of secreted high-molecular weight compounds such as lipopeptides (Berry, 2010; Geudens and Martins, 2018; Mejri et al., 2018) in a high throughput manner. However, lower throughput microscopic image analysis to assess fungal blastospore density in response to confrontation will facilitate the determination of infraspecific suppressive activity of pseudomonads.

The 11 *Pseudomonas* isolates screened for antagonism of genetically diverse *Z. tritici* isolates were analysed bioinformatically to obtain predictions of secondary metabolite BGCs from genome assemblies. This analysis indicated that known antifungal secondary metabolite BGCs, such as genes for 2,4-DAPG production, were implicated in fungal suppression *in vitro*. However, we also found that for *P. mandelii* subgroup isolate BFB20, a mechanism of antagonism through the production of known secreted antifungal secondary metabolites could not be inferred from predicted BGC similarity to MIBiG reference BGCs. Additionally, the genome assembly of *P. mandelii* isolate BFR20 only contained one predicted BGC found to be significantly similar to the MiBIG reference BGC for Pf-5 pyoverdine (BGC0000413), though BGCs in this GCF were also predicted from non-antagonistic strains Roth17, Roth40, Roth97, and Roth105. The observed antagonism phenotype of these strains, BFB20 and BFR20, could therefore be due to the production of potentially novel secondary metabolites, encoded by BGCs dissimilar to reference BGCs from the MIBiG repository. As such these strains, and predicted BGCs within GCFs deriving solely from antagonistic isolates, namely GCF_00030, GCF_00023, and GCF_00028 (all predicted from BFR20), GCF_02559 (predicted from BFR20 and Roth112), and GCF_00016 (predicted from BFB20, should be prioritised in future work for further characterisation and the potential discovery of new antifungal molecules. These findings are consistent with s with previous studies that found many predicted secondary metabolite BGCs from *Pseudomonas* type strains were significantly dissimilar to known references BGCs from the MiBIG repository, and therefore potentially encode novel secondary metabolites (Saati-Santamaría et al., 2022). Characterisation of predicted BGCs with significant similarity to reference BGCs encoding 2,4-DAPG production in isolates Roth107 and Roth112 (*P. corrugata* subgroup) will determine whether the observed *in vitro* antagonism phenotype is solely due to 2,4-DAPG production, or if other predicted BGCs encoding unknown secondary metabolite products contribute to the phenotype. However, our results showed that the *in vitro* antagonism of isolate Roth82 (*P. corrugata* subgroup) appeared to be solely due to 2,4-DAPG production.

We found that three of the antagonistic bacteria were bioinformatically predicted to have the potential to biosynthesise the antifungal polyketide 2,4-DAPG. Production of 2,4-DAPG has been described from two *Pseudomonas* species subgroups, *P. corrugata* and *P. protegens* (Abbas et al., 2002; Almario et al., 2017; Frapolli et al., 2007; Hansen et al., 2021; Raaijmakers and Weller, 1998), and more recently in the genus *Chromobacterium* (Johnson et al., 2023). This corroborates with the putative classification of Roth82, and Roth107, and Roth112 as within the *P. corrugata* subgroup based on taxonomic inference within this study, and the presence of a predicted BGC with high sequence similarity to 2,4-DAPG reference BGCs (Abbas et al., 2002; Bangera and Thomashow, 1999; Hansen et al., 2021; Raaijmakers and Weller, 1998). Predicted BGCs with a high sequence similarity to those encoding the biosynthesis of pyoverdines (Hartney et al., 2013) and pseudomonine (Mercado-Blanco et al., 2001) were also predicted from the genome assemblies of *Pseudomonas* isolates within this study but were either found in an equal number of *Z. tritici* antagonistic and non-antagonistic strains (pyoverdine), or solely from *Z. tritici* non-antagonistic isolates (pseudomonine). Whilst these BGCs were not taken forward for further characterisation in respect to the *Z. tritici* antagonism phenotype, it is possible that diversity in the expression profiles of these BGCs exist between the *Pseudomonas* isolates, or that the siderophore products of these BGCs differ between isolates. For this reason, predicted BGC presence/absence was only used to enable prioritisation of BGCs associated with *Z. tritici* antagonism phenotypes, rather than provide definitive identifications of similarity. Additionally, whilst these predicted BGCs did not appear to be strongly associated with the *Z. tritici* blastospore antagonism phenotype, they could be implicated in antagonistic interactions with other fungi, or with *Z. tritici* under different environmental conditions or contexts. For example, differing carbon sources have previously been shown to influence 2,4-DAPG production by *Pseudomonas* strain F113 (Shanahan et al., 1992). This highlights a strength of a phenotype guided-genome mining for secondary metabolites involved in microbial interactions within simplified experimental contexts.

As a proof of concept, we generated a 2,4-DAPG mutant, by disrupting the core biosynthetic enzyme PhlD. This enzyme is responsible for the production of monoacetylphloroglucinol (MAPG), that undergoes subsequent acetylation to produce 2,4-DAPG (Bangera and Thomashow, 1999). This resulted in a substantial reduction of 2,4-DAPG production in Roth82Δ*phlD*, and a loss of the visible zone of inhibition in the *Z. tritici* blastospore antagonism assay, confirming the validity of our approach to determine mode of action of known antifungal BGCs. It will be interesting to use this approach to quantify the relative contribution of predicted secondary metabolite BGCs to the *Z. tritici* antagonism phenotype where multiple antifungal BGCs are predicted in a strain. For bacterial isolates where mechanisms of antagonism cannot be bioinformatically predicted, analytical chemistry of isolate extracts can facilitate the identification of molecules implicated in fungal suppression. This, when coupled with other approaches such as the creation and screening of transposon mutant libraries can facilitate the identification of previously unknown antifungal molecules. Similarly, the *in vitro Z. tritici* blastospore antagonism assay could be adapted to screen fractions and/or compounds of interest and could be used in a bioassay-guided fractionation approach.

The efficacy of biological control agents added to environmental settings is often unpredictable (Collinge et al., 2022). Our approach of identifying molecules responsible for the suppression of *Z. tritici* does not rely on phyllosphere competence and/or production of antifungal secondary metabolites *in situ* on the plant surface. The method described can be used to efficiently and cost-effectively mine bacterial strain collections for novel antifungal secondary metabolites that are effective against *Z. tritici* in an *in vitro* setting. Furthermore, the described *Pseudomonas*-*Zymoseptoria* interaction system could be further exploited as a potential ‘stepping-stone’ for the identification of broad spectrum antifungal secondary metabolites or compounds with activity against other fungal plant pathogens, that are less amenable to high-throughput *in vitro* screening, as well as medically important fungi such as *Candida* and *Aspergillus* spp.

### Experimental procedures

#### *Pseudomonas* isolate collection

The library of 534 *Pseudomonas* wheat root associated isolates utilised in this study were generated previously at Rothamsted Research in Harpenden, UK (Mauchline et al., 2015; Ruscoe et al., 2021) and maintained as glycerol stocks in the Rothamsted microbial culture collection at −80 °C.

#### Zymoseptoria tritici isolate collection

Twelve genetically diverse *Z. tritici* isolates utilised in this study were isolated from across Europe in 2016 as part of a previous study(Chen et al., 2023), with the exception of *Z. tritici* isolate IPO323, the reference strain first isolated in the Netherlands in 1984 (Kema and Van Silfhout, 1997).

#### Culturing of Pseudomonads and *Z. tritici*

*Zymoseptoria tritici* blastospores were cultured by spreading 30 μl of defrosted glycerol stock on potato dextrose agar (PDA, Formedium) plates, before inverting and placing in darkness at 16 °C for 5 days. Blastospores were harvested using a sterile loop after 5 days of growth. *Pseudomonas* isolates were cultured from glycerol stocks, by inoculating 10 ml of LB Miller Broth (Formedium), with a chip (approximately 5 μl volume) of frozen glycerol stock. The inoculated broth was placed in a shaking incubator at 180 rpm and 25 °C overnight (18 hours).

#### *Zymoseptoria tritici* blastospore antagonism assay

To qualitatively assess *Z. tritici* antagonism, a 20 μl loop of harvested *Z. tritici* blastospores was vortexed with 5 ml of distilled water, resulting in the formation of a suspension of approximately 1×10 blastospores per ml (Chen et al., 2023). Next, 800 μl of spore suspension was added to a 120 mm x 120 mm Petri dish plates containing 50 ml PDA (Formedium). Plates were tilted to ensure a confluent inoculation of the blastospore suspension across the entire plate before excess spore suspension was recovered. Plates were air-dried for 20 minutes in a laminar flow hood, before 1 μl of 0.6 OD_600_ of a given *Pseudomonas* overnight culture was applied. A total of up to 48 *Pseudomonas* isolates were arrayed per plate in triplicate. Plates were sealed with Parafilm (Amcor) and inverted, before incubation in darkness at 16 °C for 12 days, prior to assessment for fungal growth inhibition. Images were taken after 12 days from a standardised height of 78.5 cm, using a Nikon D80 digital camera with a Sigma 70 mm F2.8 DG Macro lens (settings ISO 400, RAW image quality, large image size).

When performing quantitative antagonism measurements, the assay was conducted as described above with increased spacing; 11 *Pseudomonas* isolates and an overnight LB Miller Broth (Formedium) negative control were included per plate and four replicate plates were prepared in a factorial design. Images of confrontation plates were used to measure fungal growth inhibition as described above. Measurements were taken from the edge of the bacterial colony to the edge of the zone of clearing of *Z. tritici* blastospores, in a radial manner from the centre of the bacterial colony, using FIJI/ImageJ (version 2.1.0/1.53c) to quantify the number of pixels (2253 pixels = 1 mm). A total of 3 measurements were taken for each bacterial colony on the quantitative *Z. tritici* blastospore confrontation assay plate, using a randomly generated angle of measurement from the colony centre to the edge of the zone of clearing for each measurement. Figure 1a and Figure 2a were created using BioRender.

#### Genome sequencing of *Pseudomonas* isolates

Genome sequencing was provided by MicrobesNG (http://www.microbesng.com). Draft genome assemblies for 11 *Pseudomonas* isolates were generated using 250bp paired end Illumina MiSeq. Sequences were passed through their bioinformatic pipeline consisting of Kraken (Wood and Salzberg, 2014), BWA mem (Li, 2013), before genomes were assembled *de novo* using SPAdes (Prjibelski et al., 2020) and annotated using Prodigal (Hyatt et al., 2010).

### BUSCO (Benchmarking Universal Single-Copy Orthologs) analysis

BUSCO analysis of genomes was carried out on a local server using BUSCO version 5.2.2 (Manni et al., 2021), using the bacterial odb10 reference single copy orthologous genes dataset for bacterial species to assess genome assemblies for bacterial contamination, and the Pseudomonadales odb10 reference dataset to assess genome assembly completeness.

### Phylogenetic tree construction

A phylogenetic tree was constructed using Automated Multi-Locus Species Tree (autoMLST), based on a concatenation of 100 common genes (Table S3) found across the genome assemblies of the 11 *Pseudomonas* isolates within this study, and the closest 50 reference genome assemblies from RefSeq (Alanjary et al., 2019). Annotations to the phylogenetic tree were added using ITOL (Letunic and Bork, 2019). Clades within the phylogenetic tree were annotated, to indicate the *Pseudomonas* subgroups present as defined by Hesse *et al*. 2018 (Hesse et al., 2018). Type strain *Pseudomonas* genomes were identified using The List of Prokaryotic names with Standing in Nomenclature (LPSN) (https://lpsn.dsmz.de/) accessed on the 24th January 2025.

### Secondary metabolite biosynthetic gene cluster prediction and sequence similarity network (SSN) generation

Secondary metabolite biosynthetic gene clusters were predicted using antiSMASH 7.0.0 on a local server for 5 antagonistic and 6 non-antagonistic *Pseudomonas* genome sequences. Sequence similarity networks (SSN) were constructed from the antiSMASH 7.0.0 output from the 11 antagonistic *Pseudomonas* genome sequences, using BiG-SCAPE version 1.1.5 (Navarro-Muñoz et al., 2020) and Pfam 35 (Mistry et al., 2021) on a local server. A cut-off value of 0.3 was used, and 1,808 reference sequences from the MIBiG database were included in the construction of Gene Cluster Family (GCF) networks based on sequence similarity, with visualisations of SSNs generated and edited in Cytoscape 3.10.1 (Shannon et al., 2003). Regions containing predicted BGCs that formed a SSN with MIBiG reference BGCs for 2,4-diacetylphloroglucinol (BGC0000280 & BGC0000281) were aligned using clinker version 0.0.23(Gilchrist and Chooi, 2021), before trimming to remove sequence upstream and downstream of the predicted BGCs. Colour schemes of plots were modified using Inkscape version 1.3 (Available from: https://inkscape.org).

### Construction of *phlD* mutant and complement

To generate Roth82Δ*phlD* an internal fragment (333 bp) of the *phlD* gene was amplified by PCR using primers (pKNOCK*PhlD*3_Fw: GTCGGATCCACATCGTGCACCGGTTTCAT and pKNOCK*PhlD*3_Rv: GTCGGTACCGTAAGACCCGGTATTGGCG), verified by DNA sequencing and cloned as a *Bam*HI-*Kpn*I fragment into corresponding sites of pKNOCK-Km (Alexeyev, 1999), resulting in pKNOCK*phlD*. This plasmid construct was delivered to *Pseudomonas* isolate Roth82 by electroporation, and transformants selected after appropriate antibiotic selection. The mutant Roth82Δ*phlD* was confirmed by colony PCR using primers (pKnockPhlD2_Fw: ATGATGCCCTCGCTGAC, pKnockPhlD2_Rv: CCGGGTTCCAAGTCCAGT, pKNOCKcheckFw: GGACAACAAGCCAGGGATGTA and pKNOCKcheckRv: ATGTAAGCCCACTGCAAGCTA) and by Sanger sequencing at Eurofins (Germany) (Figure S6).

### Chemical extraction of secondary metabolites from *Z. tritici* antagonism assay

A 10 mm diameter section of agar surrounding the *Pseudomonas* colony edge was excised from wild-type Roth82 and mutant Roth82*phlD* grown with and without *Z. tritici* confrontation on PDA plates. Chemical extracts were prepared by macerating agar sections in 1 ml HPLC grade MeOH, with resulting solutions centrifuged at >13,000 rpm for 10 minutes. Supernatant was collected for analysis by liquid chromatography-mass spectrometry (LC-MS).

### Chemical analysis of secreted secondary metabolites produced by *Pseudomonas* isolates

UPLC was conducted on agar extracts using an ACQUITY UPLC® system equipped with an AQUITY BEH C18 (1.7 µm, 2.1 x 50 mm) analytical column (Waters, USA). Extracts were eluted over 38 minutes from 5% to 95% organic (UPLC grade water + 0.1% formic acid and MeOH + 0.1% formic acid) at a flow rate of 0.21 mL min. The UPLC system was coupled with Quantitative Time-of-Flight mass spectrometer SYNAPT G2-Si (Waters, USA), with electrospray ion source run in negative and positive ionisation mode and data acquired over 50-1200 m/z range. The lock mass solution consisted of Leucine Enkephalin (200 pg/µL). The lockspray was acquired during UPLC-MS acquisition and corrected for ([M+H] m/z 556.2771 & [M-H] m/z 554.2615). Instrument control and data acquisition was carried out in MassLynx (Waters, USA). Data was converted to mzML format using Waters2mzML V1.2.0 (available at https://github.com/AnP311/Waters2mzML), before Extracted Ion Chromatograms (EICs) were generated using mzmine 4.3.0(Schmid et al., 2023).

### Statistical analyses

Analysis of Variance (ANOVA) was used to determine significance difference in the size of zones of inhibition. Zone of inhibition data of *Z. tritici* isolates in confrontation with *Pseudomonas* isolate BFR20 were log transformed, to fulfil the requirement of analysis of variance (ANOVA) for data to be approximately normally distributed. Zone of inhibition data for all other antagonistic *Pseudomonas* isolates were found to be approximately normally distributed without transformation. Sidak’s multiple comparison post-hoc test was used to examine pairwise data comparisons between *Z. tritici* genotypes at the 5% level of significance. All statistical analyses were carried out in RStudio (2023.03.0 Build 386, R version 4.2.3), using packages ggplot2 (3.4.4) (Wickham, 2016), dplyr (1.1.4) (Wickham et al., 2023), ggpubr (0.6.0) (Kassambara, 2023), emmeans (1.10.0) (Lenth, 2024), multcomp (1.4-25) (Hothorn et al., 2008) and svglite (2.1.3) (Wickham et al., 2024).

## Acknowledgements

The authors would like to sincerely thank Dr. Vittorio Tracanna and Prof. Marnix Medema for providing bioinformatics training that was essential for this study during a placement hosting G.L. at Wageningen University.

The authors acknowledge funding from UK Biotechnology and Biological Sciences Research Council (BBSRC) Nottingham-Rothamsted Doctoral Training Partnership (grant number BB/M008770/1), and the Lawes Agricultural Trust. Rothamsted Science Initiatives Catalyst Award scheme grant ‘Microbial natural product discovery pipeline for next generation fungicides’

Rothamsted Research receives strategic funding from the Biotechnology and Biological Sciences Research Council of the United Kingdom. We acknowledge support from the Institute Strategic Programmes: Growing Health (BB/X010953/1; BBS/E/RH/230003A and BBS/E/RH/230003B) and Delivering Sustainable Wheat (BB/X011003/1 and BBS/E/RH/230001B), as well as previous Institute Strategic Programmes: Designing Future Wheat (BBS/E/C000I0250), Soil 2 Nutrition (BBS/E/C/000I0310), and Smart Crop Protection (SCP, BBS/OS/CP/000001), funded through BBSRC’s Industry Strategy Challenge Fund.

## Data Availability statement

The *Pseudomonas* genome assemblies analysed in this study [are/will be available following publication] from the NCBI Bioproject database (accession: PRJNA1164014) as BioSamples SAMN43882061-SAMN43882071. All scripts and processed datasets associated with this study will be made publicly available, with Datasets associated with this project will be made available under CC-BY 4.0 license in the Rothamsted Repository, alongside Python and R scripts for the analysis of the data and generation of figures available under the Apache 2– 0 license in a Github repository (https://github.com/iamgeorgelund/PseudomonasZymoseptoria), mirrored in a permanent Zenodo repository.

## Supporting Information legends

**Figure S1. A bar chart indicating the status of 124 BUSCO genes from the Bacteria odb10 dataset of the *Pseudomonas* genome assemblies utilised in this study.** All genome assemblies were found to show high completeness and low levels of contamination using the general bacterial dataset.

**Figure S2. A bar chart indicating the status of 782 BUSCO genes from the Pseudomonadales odb10 dataset of the *Pseudomonas* genome assemblies utilised in this study.** All genome assemblies were found to show high completeness and low levels of contamination using the Pseudomonadales dataset.

**Figure S3. Summary of predicted secondary metabolite biosynthetic gene cluster classes.** A bar chart summarising the number of predicted secondary metabolite biosynthetic gene clusters and their biosynthetic classes from the genome assemblies of each *Pseudomonas* strain tested within this study.

**Figure S4. BiG-SCAPE sequence similarity networks of predicted BGCs from the 11 *Pseudomonas* isolate genome assemblies.** Sequence similarity networks of predicted BGCs, with nodes coloured according to the isolate genome assembly from which they were predicted. Connected nodes indicate similarity at the 0.3 threshold.

**Figure S5. Phylogenetic distribution of predicted secondary metabolite BGCs and*Z. Tritici* blastopore antagonism phenotypes of *Pseudomonas* isolates used in this study.** Phylogenetic analysis with known *Pseudomonas* strains, using autoMLST, putatively identifies 11 *Pseudomonas* strains used within this study within the *P. fluorescens*, *P. corrugata*, *P. koreensis*, *P. jessenii*, and *P. mandelii* subgroups of the *P. fluorescens* species complex. The presence and absence of all predicted BGCs that form Gene Cluster Family sequence similarity networks (GCFs) are indicated, with those alongside qualitative *Z. tritici* blastospore antagonism phenotype against *Z. tritici* strain IPO323. Type strain *Pseudomonas* genomes identified according to LPSN (accessed 24 January 2025) are denoted by a superscript ‘T’.

**Figure S6. Agarose gel electrophoresis of PCR (using external primers from the insertion region, pKnockPhlD2_Fw – pKnockPhlD2_Rv) for verification of *phlD* gene knockout.** Roth82 PCR product size is 527bp, Roth82*ΔphlD* shows the insertion of pKnock-Km vector (2098bp).

**Table S1. 52 *Pseudomonas* isolates from the culture collection of 534 identified as antagonistic in qualitative antagonism assay against *Z. tritici* isolate IPO323.**

**Table S2. Qualitative *Z. tritici* antagonism phenotype of *Pseudomonas* isolates used in the quantitative antagonism assay against 12 diverse *Z. tritici* isolates.**

**Table S3. Genes utilised in the construction of the phylogenetic tree using autoMLST.**

