## Supporting Information for "Development of a bioassay guided genome mining approach for antifungal natural product discovery from pseudomonads"

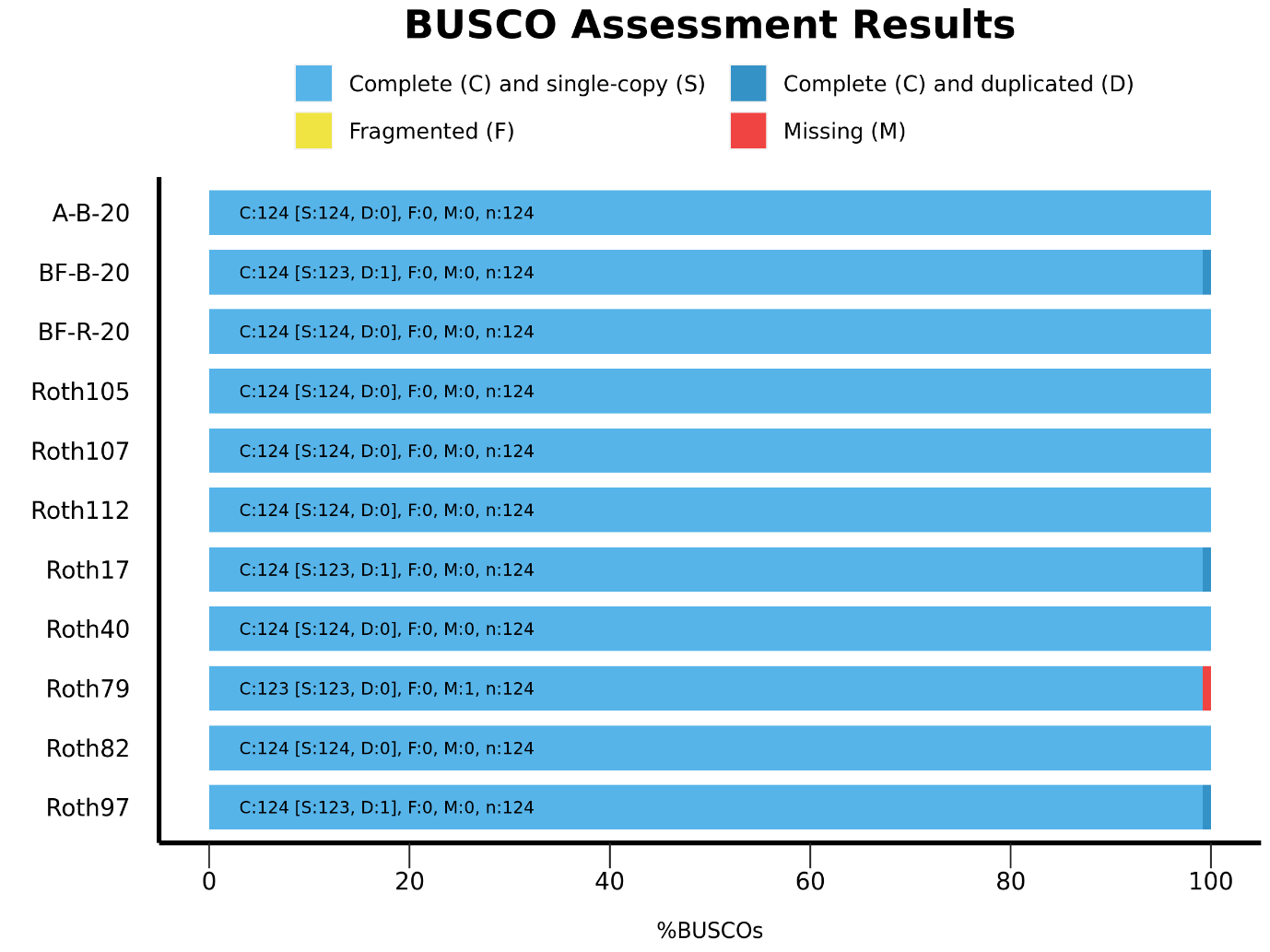

**Figure S1. A bar chart indicating the status of 124 BUSCO genes from the Bacteria odb10 dataset of the *Pseudomonas* genome assemblies utilised in this study.** All genome assemblies were found to show high completeness and low levels of contamination using the general bacterial dataset.

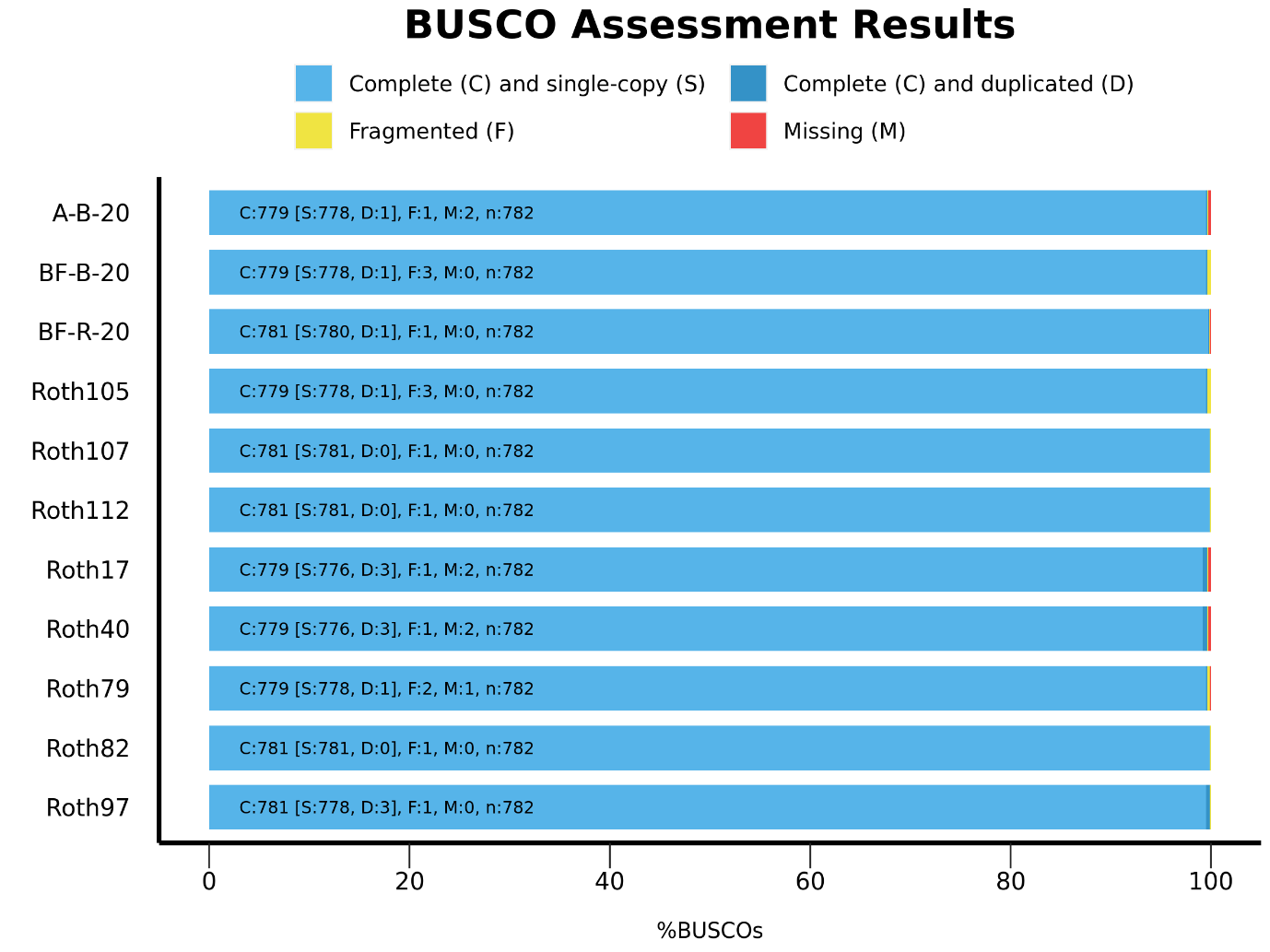

**Figure S2. A bar chart indicating the status of 782 BUSCO genes from the Pseudomonadales odb10 dataset of the *Pseudomonas* genome assemblies utilised in this study.** All genome assemblies were found to show high completeness and low levels of contamination using the Pseudomonadales dataset.

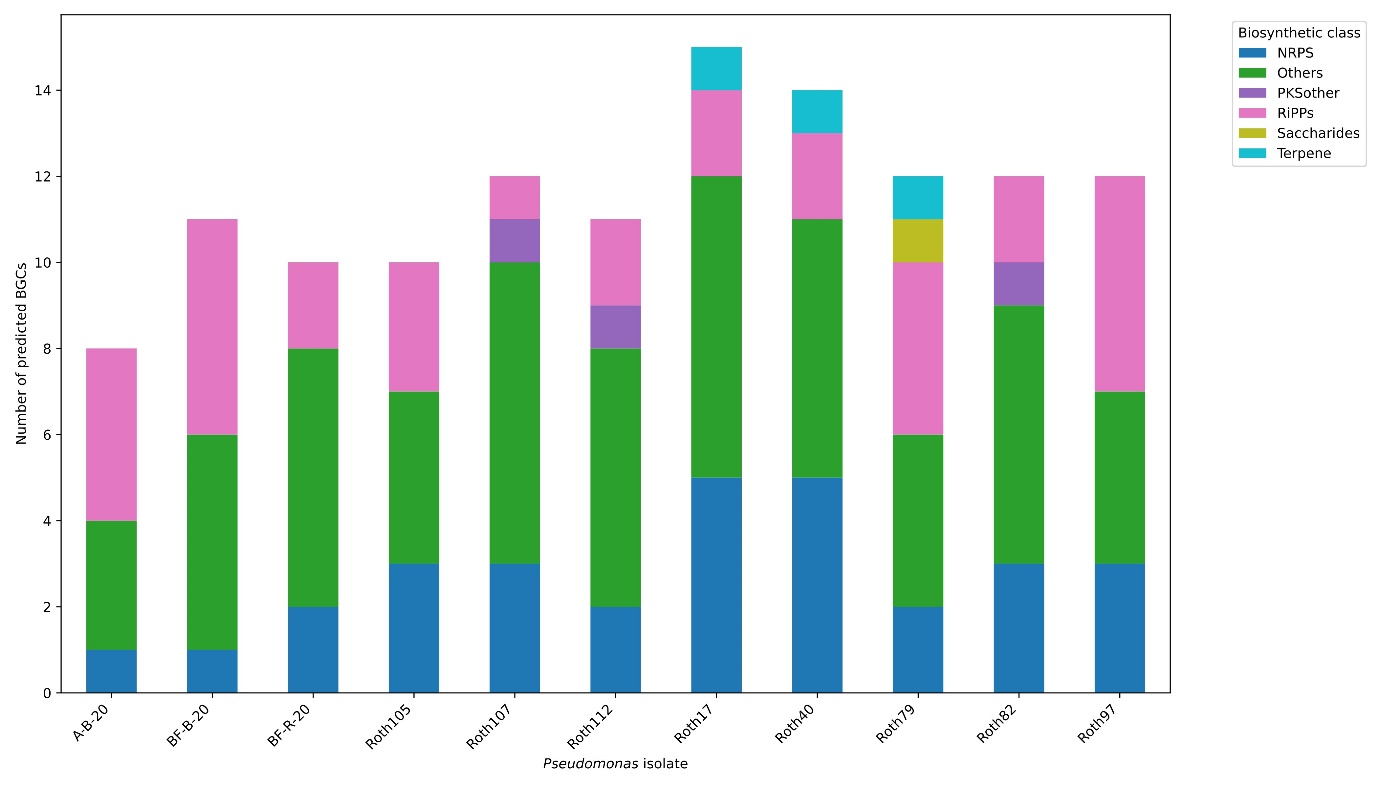

**Figure S3. Summary of predicted secondary metabolite biosynthetic gene cluster classes.** A bar chart summarising the number of predicted secondary metabolite biosynthetic gene clusters and their biosynthetic classes from the genome assemblies of each *Pseudomonas* strain tested within this study.

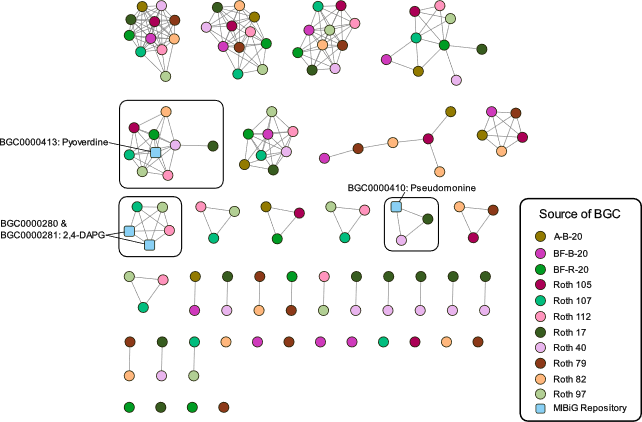

**Figure S4. BiG-SCAPE sequence similarity networks of predicted BGCs from the 11 *Pseudomonas* isolate genome assemblies.** Sequence similarity networks of predicted BGCs, with nodes coloured according to the isolate genome assembly from which they were predicted. Connected nodes indicate similarity at the 0.3 threshold.

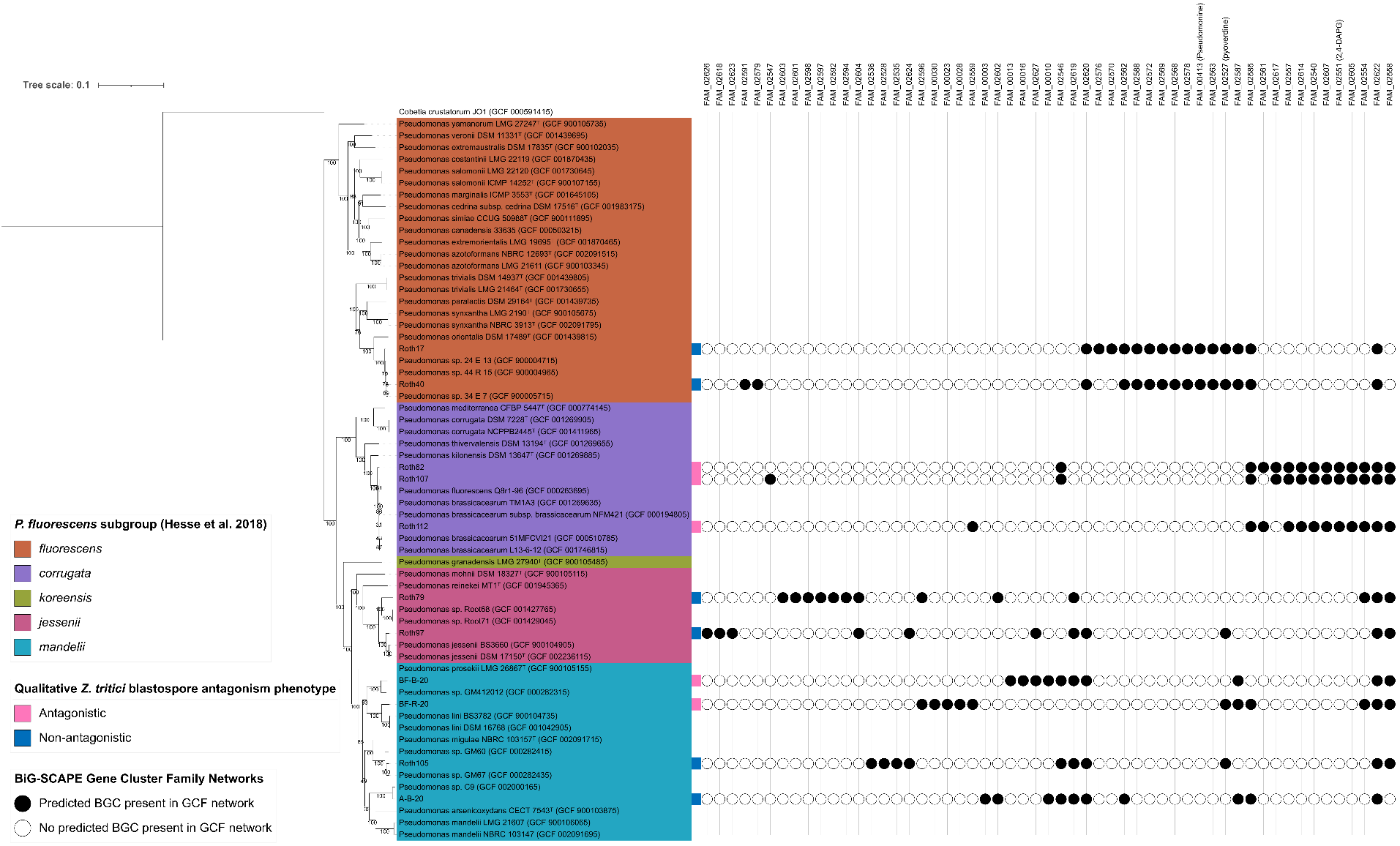

**Figure S5. Phylogenetic distribution of predicted secondary metabolite BGCs and *Z. tritici* blastopore antagonism phenotypes of *Pseudomonas* isolates used in this study.** Phylogenetic analysis with known *Pseudomonas* strains, using autoMLST, putatively identifies 11 *Pseudomonas* strains used within this study within the *P. fluorescens*, *P. corrugata*, *P. koreensis*, *P. jessenii*, and *P. mandelii* subgroups of the *P. fluorescens* species complex. The presence and absence of all predicted BGCs that form Gene Cluster Family sequence similarity networks (GCFs) are indicated, with those alongside qualitative *Z. tritici* blastospore antagonism phenotype against *Z. tritici* strain IPO323. Type strain *Pseudomonas* genomes identified according to LPSN (accessed 24^th^ January 2025) are denoted by a superscript ‘T’.

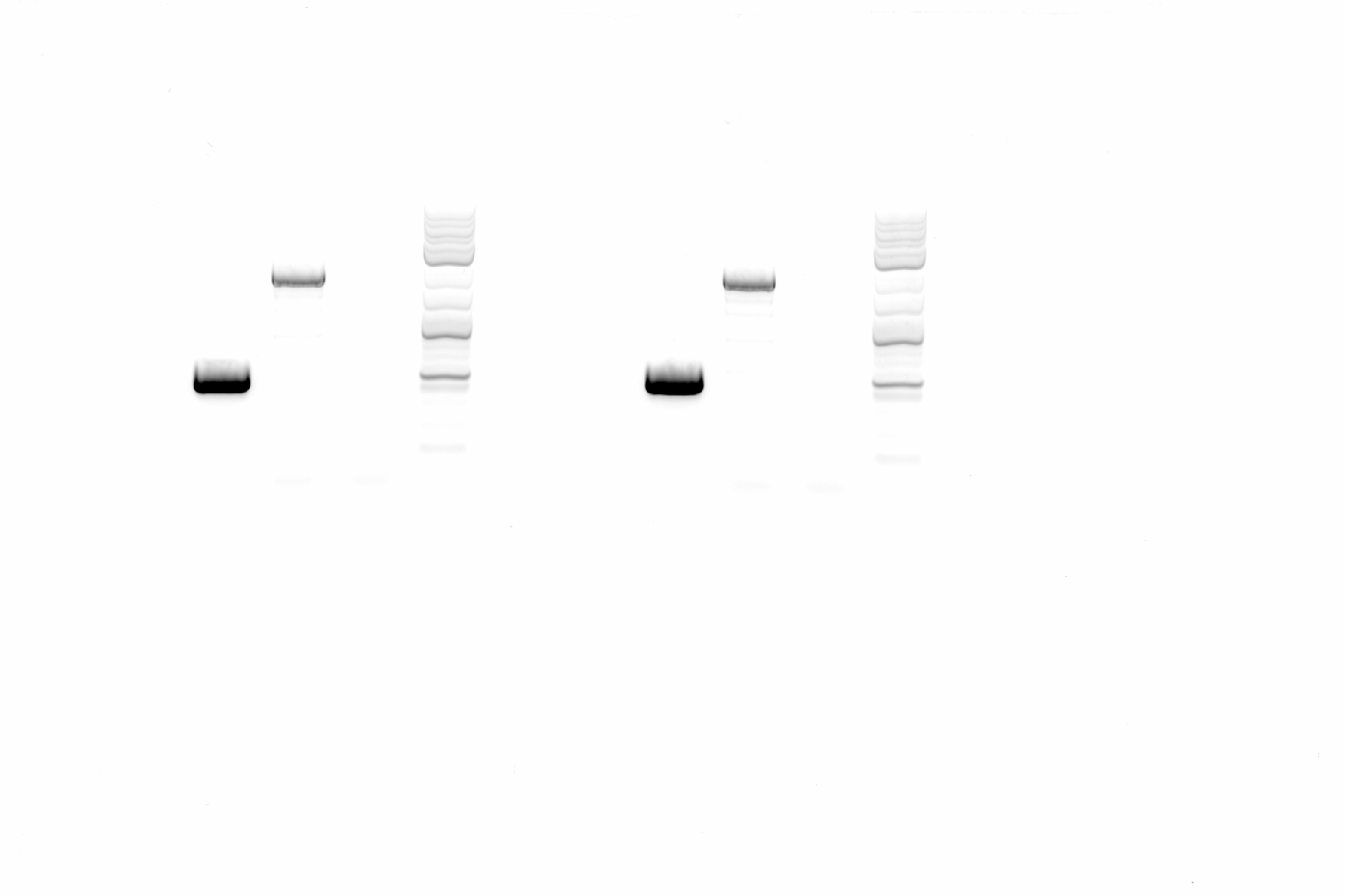

**Roth82**

**Roth82∆*phlD***

**water**

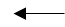

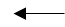

500bp

2000bp

3000bp

**Figure S6.** **Agarose gel electrophoresis of PCR (using external primers from the insertion region, pKnockPhlD2_Fw – pKnockPhlD2_Rv) for verification of *phlD* gene knockout.** Roth82 PCR product size is 527bp, Roth82*∆phlD* shows the insertion of pKnock-Km vector (2098bp).

**Table S1. 52 *Pseudomonas* isolates from the culture collection of 534 identified as antagonistic in qualitative antagonism assay against *Z. tritici* isolate IPO323.**

| ***Pseudomonas* isolate** | ***Z. tritici* qualitative blastospore antagonism phenotype** |
| --- | --- |
| 1E_1 | antagonistic |
| 1E_2 | antagonistic |
| 1R_3 | antagonistic |
| 22E_6 | antagonistic |
| 24E_10 | antagonistic |
| 24R_5 | antagonistic |
| 28E_14 | antagonistic |
| 28E_15 | antagonistic |
| 28R_11 | antagonistic |
| 28R_2 | antagonistic |
| 28R_7 | antagonistic |
| 30R_14 | antagonistic |
| 30R_16 | antagonistic |
| 30R_2 | antagonistic |
| 30R_8 | antagonistic |
| 31E_11 | antagonistic |
| 31R_14 | antagonistic |
| 35E_3 | antagonistic |
| 37R_16 | antagonistic |
| 37R_6 | antagonistic |
| 37R_7 | antagonistic |
| 37R_8 | antagonistic |
| 58E_12 | antagonistic |
| 58E_13 | antagonistic |
| 58R_1 | antagonistic |
| 58R_2 | antagonistic |
| 8E_1 | antagonistic |
| BF-B-20 | antagonistic |
| BF-R-20 | antagonistic |
| Roth103 | antagonistic |
| Roth104 | antagonistic |
| Roth107 | antagonistic |
| Roth111 | antagonistic |
| Roth112 | antagonistic |
| Roth12 | antagonistic |
| Roth16 | antagonistic |
| Roth20 | antagonistic |
| Roth21 | antagonistic |
| Roth26 | antagonistic |
| Roth4 | antagonistic |
| Roth40 | antagonistic |
| Roth54 | antagonistic |
| Roth56 | antagonistic |
| Roth58 | antagonistic |
| Roth64 | antagonistic |
| Roth65 | antagonistic |
| Roth70 | antagonistic |
| Roth73 | antagonistic |
| Roth79 | antagonistic |
| Roth81 | antagonistic |
| Roth82 | antagonistic |
| Roth83 | antagonistic |
| Roth97 | antagonistic |

**Table S2. Qualitative *Z. tritici* antagonism phenotype of *Pseudomonas* isolates used in the quantitative antagonism assay against 12 diverse *Z. tritici* isolates.**

| ***Pseudomonas* isolate** | ***Z. tritici* qualitative blastospore antagonism phenotype** |
| --- | --- |
| A-B-20 | non-antagonistic |
| BF-B-20 | antagonistic |
| BF-R-20 | antagonistic |
| Roth17 | non-antagonistic |
| Roth40 | non-antagonistic |
| Roth79 | non-antagonistic |
| Roth82 | antagonistic |
| Roth97 | non-antagonistic |
| Roth105 | non-antagonistic |
| Roth107 | antagonistic |
| Roth112 | antagonistic |

**Table S3. Genes utilised in the construction of the phylogenetic tree using autoMLST.**

TIGR01067

TIGR01050

TIGR01164

TIGR01032

TIGR01029

TIGR00493

TIGR00138

TIGR01966

TIGR03635

TIGR03284

TIGR00952

TIGR02012

TIGR01169

TIGR01011

TIGR00967

TIGR00928

TIGR03723

TIGR00612

TIGR01302

TIGR00690

TIGR00019

TIGR00362

TIGR01009

TIGR00962

TIGR00628

TIGR00521

TIGR01039

TIGR03632

TIGR00482

TIGR01850

TIGR00132

TIGR01021

TIGR00048

TIGR00713

TIGR00855

TIGR02970

TIGR00563

TIGR02075

TIGR01394

TIGR01207

TIGR00338

TIGR00263

TIGR00008

TIGR00067

TIGR02673

TIGR00382

TIGR02027

TIGR00651

TIGR00337

TIGR00767

TIGR00088

TIGR00418

TIGR03346

TIGR01792

TIGR00190

TIGR00963

TIGR01171

TIGR00635

TIGR00060

TIGR00431

TIGR00243

TIGR02225

TIGR00228

TIGR00012

TIGR00062

TIGR02274

TIGR03631

TIGR01059

TIGR00755

TIGR01163

TIGR00184

TIGR00615

TIGR00065

TIGR00119

TIGR00033

TIGR00708

TIGR01049

TIGR00165

TIGR01632

TIGR03594

TIGR00355

TIGR00168

TIGR00508

TIGR00036

TIGR01044

TIGR00055

TIGR00981

TIGR01024

TIGR01083

TIGR00959

TIGR00430

TIGR00118

TIGR00459

TIGR02729

TIGR00499

TIGR01816

TIGR02013

TIGR00580

TIGR02386

Table S4. One-way Analysis of Variance of the radius of zones of inhibition produced by *Pseudomonas* isolate BFR20 in confrontation with genetically diverse *Z. tritici* isolates in the blastospore antagonism assay *in vitro*.

Measurements were log transformed to achieve approximate normality.

|  | Df | Sum Sq | Mean Sq | *F* value | Pr(>*F*) |  |
| --- | --- | --- | --- | --- | --- | --- |
| *Z. tritici* genotype | 11 | 45.198 | 4.109 | 11.476 | 9.423E-09 | *** |
| Residuals | 36 | 12.889 | 0.358 |  |  |  |

Significance codes: 0 ‘***’ 0.001 ‘**’ 0.01 ‘*’ 0.05 ‘.’ 0.1 ‘ ’ 1

Table S5. Tukey’s test for multiple comparisons at the 5% level of significance, comparing zones of inhibition produced by *Pseudomonas* isolate BFR20 in confrontation with different *Z. tritici* genotypes *in vitro*.

Measurements were log transformed to achieve approximate normality.

| Comparison | Mean Difference | Lower Bound | Upper Bound | Padj |
| --- | --- | --- | --- | --- |
| L951-IPO323 | 3.3892755 | 1.91250399 | 4.86604702 | 0.0000001 |
| Zt10-IPO323 | 2.1189732 | 0.64220166 | 3.59574469 | 0.0007863 |
| Zt114-IPO323 | 2.3035831 | 0.82681158 | 3.78035461 | 0.0002151 |
| Zt116-IPO323 | 3.6951467 | 2.21837519 | 5.17191822 | 0 |
| Zt118-IPO323 | 2.9357476 | 1.45897613 | 4.41251916 | 0.0000024 |
| Zt120-IPO323 | 3.8555348 | 2.37876325 | 5.33230628 | 0 |
| Zt36-IPO323 | 1.9692034 | 0.49243186 | 3.44597489 | 0.0022042 |
| Zt55-IPO323 | 2.5081041 | 1.0313326 | 3.98487563 | 0.0000502 |
| Zt74-IPO323 | 3.0945381 | 1.61776662 | 4.57130965 | 0.0000008 |
| Zt80-IPO323 | 2.7117675 | 1.23499602 | 4.18853905 | 0.0000118 |
| Zt92-IPO323 | 2.8210343 | 1.34426279 | 4.29780583 | 0.0000054 |
| Zt10-L951 | -1.2703023 | -2.74707384 | 0.20646919 | 0.1494898 |
| Zt114-L951 | -1.0856924 | -2.56246393 | 0.39107911 | 0.3348054 |
| Zt116-L951 | 0.3058712 | -1.17090032 | 1.78264272 | 0.9998325 |
| Zt118-L951 | -0.4535279 | -1.93029938 | 1.02324365 | 0.9941461 |
| Zt120-L951 | 0.4662593 | -1.01051225 | 1.94303078 | 0.9926561 |
| Zt36-L951 | -1.4200721 | -2.89684365 | 0.05669938 | 0.0686984 |
| Zt55-L951 | -0.8811714 | -2.35794291 | 0.59560013 | 0.6372617 |
| Zt74-L951 | -0.2947374 | -1.77150888 | 1.18203415 | 0.999883 |
| Zt80-L951 | -0.677508 | -2.15427948 | 0.79926355 | 0.8974541 |
| Zt92-L951 | -0.5682412 | -2.04501271 | 0.90853032 | 0.9669885 |
| Zt114-Zt10 | 0.1846099 | -1.2921616 | 1.66138143 | 0.999999 |
| Zt116-Zt10 | 1.5761735 | 0.09940201 | 3.05294504 | 0.027913 |
| Zt118-Zt10 | 0.8167745 | -0.65999705 | 2.29354598 | 0.7334086 |
| Zt120-Zt10 | 1.7365616 | 0.25979008 | 3.21333311 | 0.0102865 |
| Zt36-Zt10 | -0.1497698 | -1.62654132 | 1.32700171 | 0.9999999 |
| Zt55-Zt10 | 0.3891309 | -1.08764058 | 1.86590245 | 0.9984292 |
| Zt74-Zt10 | 0.975565 | -0.50120655 | 2.45233648 | 0.4909093 |
| Zt80-Zt10 | 0.5927944 | -0.88397715 | 2.06956588 | 0.9558116 |
| Zt92-Zt10 | 0.7020611 | -0.77471038 | 2.17883265 | 0.8744014 |
| Zt116-Zt114 | 1.3915636 | -0.08520791 | 2.86833513 | 0.080234 |
| Zt118-Zt114 | 0.6321645 | -0.84460697 | 2.10893606 | 0.9326965 |
| Zt120-Zt114 | 1.5519517 | 0.07518016 | 3.02872319 | 0.032264 |
| Zt36-Zt114 | -0.3343797 | -1.81115124 | 1.14239179 | 0.9996084 |
| Zt55-Zt114 | 0.204521 | -1.2722505 | 1.68129254 | 0.9999971 |
| Zt74-Zt114 | 0.790955 | -0.68581647 | 2.26772656 | 0.7692799 |
| Zt80-Zt114 | 0.4081844 | -1.06858707 | 1.88495596 | 0.9976067 |
| Zt92-Zt114 | 0.5174512 | -0.9593203 | 1.99422273 | 0.9833483 |
| Zt118-Zt116 | -0.7593991 | -2.23617058 | 0.71737245 | 0.8101745 |
| Zt120-Zt116 | 0.1603881 | -1.31638345 | 1.63715958 | 0.9999998 |
| Zt36-Zt116 | -1.7259433 | -3.20271485 | -0.24917182 | 0.0110099 |
| Zt55-Zt116 | -1.1870426 | -2.66381411 | 0.28972893 | 0.2201155 |
| Zt74-Zt116 | -0.6006086 | -2.07738008 | 0.87616295 | 0.9517499 |
| Zt80-Zt116 | -0.9833792 | -2.46015068 | 0.49339235 | 0.47905 |
| Zt92-Zt116 | -0.8741124 | -2.35088391 | 0.60265912 | 0.6481235 |
| Zt120-Zt118 | 0.9197871 | -0.55698439 | 2.39655864 | 0.5772292 |
| Zt36-Zt118 | -0.9665443 | -2.44331579 | 0.51022725 | 0.5046965 |
| Zt55-Zt118 | -0.4276435 | -1.90441504 | 1.04912799 | 0.9964248 |
| Zt74-Zt118 | 0.1587905 | -1.31798102 | 1.63556201 | 0.9999998 |
| Zt80-Zt118 | -0.2239801 | -1.70075162 | 1.25279141 | 0.9999925 |
| Zt92-Zt118 | -0.1147133 | -1.59148485 | 1.36205818 | 1 |
| Zt36-Zt120 | -1.8863314 | -3.36310291 | -0.40955988 | 0.0038541 |
| Zt55-Zt120 | -1.3474307 | -2.82420217 | 0.12934086 | 0.1013749 |
| Zt74-Zt120 | -0.7609966 | -2.23776815 | 0.71577488 | 0.8081911 |
| Zt80-Zt120 | -1.1437672 | -2.62053875 | 0.33300428 | 0.2652239 |
| Zt92-Zt120 | -1.0345005 | -2.51127197 | 0.44227106 | 0.4040227 |
| Zt55-Zt36 | 0.5389007 | -0.93787077 | 2.01567226 | 0.9774503 |
| Zt74-Zt36 | 1.1253348 | -0.35143675 | 2.60210628 | 0.2862063 |
| Zt80-Zt36 | 0.7425642 | -0.73420735 | 2.21933568 | 0.8304654 |
| Zt92-Zt36 | 0.8518309 | -0.62494058 | 2.32860245 | 0.681987 |
| Zt74-Zt55 | 0.586434 | -0.89033749 | 2.06320554 | 0.9589337 |
| Zt80-Zt55 | 0.2036634 | -1.27310809 | 1.68043494 | 0.9999972 |
| Zt92-Zt55 | 0.3129302 | -1.16384132 | 1.78970171 | 0.9997914 |
| Zt80-Zt74 | -0.3827706 | -1.85954212 | 1.09400091 | 0.9986441 |
| Zt92-Zt74 | -0.2735038 | -1.75027534 | 1.20326769 | 0.9999439 |
| Zt92-Zt80 | 0.1092668 | -1.36750474 | 1.58603829 | 1 |

Table S6. One-way Analysis of Variance of the radius of zones of inhibition produced by *Pseudomonas* isolate BFB20 in confrontation with genetically diverse *Z. tritici* isolates in the blastospore antagonism assay *in vitro*.

|  | Df | Sum Sq | Mean Sq | *F* value | Pr(>*F*) |  |
| --- | --- | --- | --- | --- | --- | --- |
| *Z. tritici* genotype | 11 | 11.846 | 1.0769 | 10.97 | 1.70E-08 | *** |
| Residuals | 36 | 3.535 | 0.0982 |  |  |  |

Significance codes: 0 ‘***’ 0.001 ‘**’ 0.01 ‘*’ 0.05 ‘.’ 0.1 ‘ ’ 1

Table S7. Tukey’s test for multiple comparisons at the 5% level of significance, comparing zones of inhibition produced by *Pseudomonas* isolate BFB20 in confrontation with different *Z. tritici* genotypes *in vitro*.

| Comparison | Mean Difference | Lower Bound | Upper Bound | Padj |
| --- | --- | --- | --- | --- |
| L951-IPO323 | 0.87613032 | 0.10274959 | 1.649511048 | 0.0153963 |
| Zt10-IPO323 | 1.27820922 | 0.50482849 | 2.051589949 | 0.0000812 |
| Zt114-IPO323 | 1.11975177 | 0.34637104 | 1.893132502 | 0.0006879 |
| Zt116-IPO323 | 0.12457004 | -0.64881069 | 0.897950764 | 0.9999862 |
| Zt118-IPO323 | 0.17792996 | -0.59545076 | 0.951310693 | 0.9995452 |
| Zt120-IPO323 | 0.67016844 | -0.10321229 | 1.443549169 | 0.1427984 |
| Zt36-IPO323 | 1.36719858 | 0.59381785 | 2.140579311 | 0.0000242 |
| Zt55-IPO323 | -0.10867021 | -0.88205094 | 0.664710516 | 0.9999966 |
| Zt74-IPO323 | 1.08843085 | 0.31505012 | 1.86181158 | 0.0010422 |
| Zt80-IPO323 | 0.40976064 | -0.36362009 | 1.183141367 | 0.78066 |
| Zt92-IPO323 | 0.83261525 | 0.05923452 | 1.605995977 | 0.0256969 |
| Zt10-L951 | 0.4020789 | -0.37130183 | 1.175459629 | 0.7996747 |
| Zt114-L951 | 0.24362145 | -0.52975927 | 1.017002183 | 0.9927912 |
| Zt116-L951 | -0.75156028 | -1.52494101 | 0.021820445 | 0.0632204 |
| Zt118-L951 | -0.69820035 | -1.47158108 | 0.075180374 | 0.1091171 |
| Zt120-L951 | -0.20596188 | -0.97934261 | 0.56741885 | 0.9982739 |
| Zt36-L951 | 0.49106826 | -0.28231247 | 1.264448991 | 0.5493102 |
| Zt55-L951 | -0.98480053 | -1.75818126 | -0.211419803 | 0.0040078 |
| Zt74-L951 | 0.21230053 | -0.5610802 | 0.985681261 | 0.9977461 |
| Zt80-L951 | -0.46636968 | -1.23975041 | 0.307011048 | 0.6227831 |
| Zt92-L951 | -0.04351507 | -0.8168958 | 0.729865658 | 1 |
| Zt114-Zt10 | -0.15845745 | -0.93183818 | 0.614923282 | 0.9998491 |
| Zt116-Zt10 | -1.15363918 | -1.92701991 | -0.380258455 | 0.0004374 |
| Zt118-Zt10 | -1.10027926 | -1.87365998 | -0.326898526 | 0.000891 |
| Zt120-Zt10 | -0.60804078 | -1.38142151 | 0.165339949 | 0.2465025 |
| Zt36-Zt10 | 0.08898936 | -0.68439137 | 0.862370091 | 0.9999996 |
| Zt55-Zt10 | -1.38687943 | -2.16026016 | -0.613498704 | 0.0000185 |
| Zt74-Zt10 | -0.18977837 | -0.9631591 | 0.58360236 | 0.9991741 |
| Zt80-Zt10 | -0.86844858 | -1.64182931 | -0.095067853 | 0.0168749 |
| Zt92-Zt10 | -0.44559397 | -1.2189747 | 0.327786757 | 0.6834417 |
| Zt116-Zt114 | -0.99518174 | -1.76856247 | -0.221801009 | 0.0035101 |
| Zt118-Zt114 | -0.94182181 | -1.71520254 | -0.16844108 | 0.0068922 |
| Zt120-Zt114 | -0.44958333 | -1.22296406 | 0.323797396 | 0.6719591 |
| Zt36-Zt114 | 0.24744681 | -0.52593392 | 1.020827537 | 0.991822 |
| Zt55-Zt114 | -1.22842199 | -2.00180271 | -0.455041257 | 0.0001596 |
| Zt74-Zt114 | -0.03132092 | -0.80470165 | 0.742059807 | 1 |
| Zt80-Zt114 | -0.70999114 | -1.48337186 | 0.063389594 | 0.0970733 |
| Zt92-Zt114 | -0.28713653 | -1.06051725 | 0.486244204 | 0.9744221 |
| Zt118-Zt116 | 0.05335993 | -0.7200208 | 0.826740658 | 1 |
| Zt120-Zt116 | 0.5455984 | -0.22778232 | 1.318979133 | 0.3937203 |
| Zt36-Zt116 | 1.24262855 | 0.46924782 | 2.016009275 | 0.0001317 |
| Zt55-Zt116 | -0.23324025 | -1.00662098 | 0.540140481 | 0.9949645 |
| Zt74-Zt116 | 0.96386082 | 0.19048009 | 1.737241545 | 0.0052268 |
| Zt80-Zt116 | 0.2851906 | -0.48819013 | 1.058571332 | 0.9756552 |
| Zt92-Zt116 | 0.70804521 | -0.06533552 | 1.481425942 | 0.0989799 |
| Zt120-Zt118 | 0.49223848 | -0.28114225 | 1.265619204 | 0.545834 |
| Zt36-Zt118 | 1.18926862 | 0.41588789 | 1.962649346 | 0.0002709 |
| Zt55-Zt118 | -0.28660018 | -1.05998091 | 0.486780552 | 0.9747666 |
| Zt74-Zt118 | 0.91050089 | 0.13712016 | 1.683881616 | 0.0101527 |
| Zt80-Zt118 | 0.23183067 | -0.54155006 | 1.005211403 | 0.9952131 |
| Zt92-Zt118 | 0.65468528 | -0.11869545 | 1.428066013 | 0.1647201 |
| Zt36-Zt120 | 0.69703014 | -0.07635059 | 1.470410871 | 0.1103778 |
| Zt55-Zt120 | -0.77883865 | -1.55221938 | -0.005457924 | 0.0471057 |
| Zt74-Zt120 | 0.41826241 | -0.35511832 | 1.19164314 | 0.7587653 |
| Zt80-Zt120 | -0.2604078 | -1.03378853 | 0.512972927 | 0.9877408 |
| Zt92-Zt120 | 0.16244681 | -0.61093392 | 0.935827537 | 0.9998082 |
| Zt55-Zt36 | -1.47586879 | -2.24924952 | -0.702488066 | 0.0000055 |
| Zt74-Zt36 | -0.27876773 | -1.05214846 | 0.494612998 | 0.9794097 |
| Zt80-Zt36 | -0.95743794 | -1.73081867 | -0.184057215 | 0.0056674 |
| Zt92-Zt36 | -0.53458333 | -1.30796406 | 0.238797395 | 0.4236466 |
| Zt74-Zt55 | 1.19710106 | 0.42372034 | 1.970481793 | 0.0002438 |
| Zt80-Zt55 | 0.51843085 | -0.25494988 | 1.29181158 | 0.4691538 |
| Zt92-Zt55 | 0.94128546 | 0.16790473 | 1.71466619 | 0.0069385 |
| Zt80-Zt74 | -0.67867021 | -1.45205094 | 0.094710516 | 0.1317931 |
| Zt92-Zt74 | -0.2558156 | -1.02919633 | 0.517565126 | 0.9893376 |
| Zt92-Zt80 | 0.42285461 | -0.35052612 | 1.196235339 | 0.7465985 |

Table S8. One-way Analysis of Variance of the radius of zones of inhibition produced by *Pseudomonas* isolate Roth82 in confrontation with genetically diverse *Z. tritici* isolates in the blastospore antagonism assay *in vitro*.

|  | Df | Sum Sq | Mean Sq | *F* value | Pr(>*F*) |  |
| --- | --- | --- | --- | --- | --- | --- |
| *Z. tritici* genotype | 11 | 20.65 | 1.8769 | 4.455 | 3.09E-04 | *** |
| Residuals | 36 | 15.16 | 0.4213 |  |  |  |

Significance codes: 0 ‘***’ 0.001 ‘**’ 0.01 ‘*’ 0.05 ‘.’ 0.1 ‘ ’ 1

Table S9. Tukey’s test for multiple comparisons at the 5% level of significance, comparing zones of inhibition produced by *Pseudomonas* isolate Roth82 in confrontation with different *Z. tritici* genotypes *in vitro*.

| Comparison | Mean Difference | Lower Bound | Upper Bound | Padj |
| --- | --- | --- | --- | --- |
| L951-IPO323 | -1.930474289 | -3.5323385 | -0.3286101 | 0.0077849 |
| Zt10-IPO323 | -1.235944148 | -2.8378084 | 0.3659201 | 0.2700704 |
| Zt114-IPO323 | -1.158492905 | -2.7603571 | 0.4433713 | 0.357921 |
| Zt116-IPO323 | -2.043147161 | -3.6450114 | -0.441283 | 0.0039252 |
| Zt118-IPO323 | -2.58317819 | -4.1850424 | -0.981314 | 0.0001238 |
| Zt120-IPO323 | -1.4358289 | -3.0376931 | 0.1660353 | 0.1145686 |
| Zt36-IPO323 | -2.016143615 | -3.6180078 | -0.4142794 | 0.004633 |
| Zt55-IPO323 | -1.930917553 | -3.5327818 | -0.3290533 | 0.0077643 |
| Zt74-IPO323 | -2.34201241 | -3.9438766 | -0.7401482 | 0.0005943 |
| Zt80-IPO323 | -2.011981383 | -3.6138456 | -0.4101172 | 0.0047525 |
| Zt92-IPO323 | -1.993377658 | -3.5952419 | -0.3915135 | 0.0053238 |
| Zt10-L951 | 0.694530141 | -0.9073341 | 2.2963943 | 0.927074 |
| Zt114-L951 | 0.771981383 | -0.8298828 | 2.3738456 | 0.8646035 |
| Zt116-L951 | -0.112672873 | -1.7145371 | 1.4891913 | 1 |
| Zt118-L951 | -0.652703902 | -2.2545681 | 0.9491603 | 0.9511446 |
| Zt120-L951 | 0.494645388 | -1.1072188 | 2.0965096 | 0.9938759 |
| Zt36-L951 | -0.085669326 | -1.6875335 | 1.5161949 | 1 |
| Zt55-L951 | -0.000443265 | -1.6023075 | 1.6014209 | 1 |
| Zt74-L951 | -0.411538121 | -2.0134023 | 1.1903261 | 0.9987476 |
| Zt80-L951 | -0.081507095 | -1.6833713 | 1.5203571 | 1 |
| Zt92-L951 | -0.062903369 | -1.6647676 | 1.5389608 | 1 |
| Zt114-Zt10 | 0.077451243 | -1.524413 | 1.6793154 | 1 |
| Zt116-Zt10 | -0.807203013 | -2.4090672 | 0.7946612 | 0.8285826 |
| Zt118-Zt10 | -1.347234042 | -2.9490982 | 0.2546302 | 0.1711614 |
| Zt120-Zt10 | -0.199884753 | -1.801749 | 1.4019795 | 0.999999 |
| Zt36-Zt10 | -0.780199467 | -2.3820637 | 0.8216647 | 0.8566073 |
| Zt55-Zt10 | -0.694973405 | -2.2968376 | 0.9068908 | 0.9267834 |
| Zt74-Zt10 | -1.106068262 | -2.7079325 | 0.4957959 | 0.4252301 |
| Zt80-Zt10 | -0.776037235 | -2.3779014 | 0.825827 | 0.860689 |
| Zt92-Zt10 | -0.75743351 | -2.3592977 | 0.8444307 | 0.8781279 |
| Zt116-Zt114 | -0.884654256 | -2.4865185 | 0.71721 | 0.7351242 |
| Zt118-Zt114 | -1.424685285 | -3.0265495 | 0.1771789 | 0.1207081 |
| Zt120-Zt114 | -0.277335995 | -1.8792002 | 1.3245282 | 0.9999713 |
| Zt36-Zt114 | -0.857650709 | -2.4595149 | 0.7442135 | 0.7696577 |
| Zt55-Zt114 | -0.772424648 | -2.3742889 | 0.8294396 | 0.8641787 |
| Zt74-Zt114 | -1.183519504 | -2.7853837 | 0.4183447 | 0.3279098 |
| Zt80-Zt114 | -0.853488478 | -2.4553527 | 0.7483757 | 0.7748113 |
| Zt92-Zt114 | -0.834884753 | -2.436749 | 0.7669795 | 0.7972338 |
| Zt118-Zt116 | -0.540031029 | -2.1418952 | 1.0618332 | 0.9876231 |
| Zt120-Zt116 | 0.607318261 | -0.9945459 | 2.2091825 | 0.9702701 |
| Zt36-Zt116 | 0.027003547 | -1.5748607 | 1.6288678 | 1 |
| Zt55-Zt116 | 0.112229608 | -1.4896346 | 1.7140938 | 1 |
| Zt74-Zt116 | -0.298865249 | -1.9007295 | 1.302999 | 0.9999396 |
| Zt80-Zt116 | 0.031165778 | -1.5706984 | 1.63303 | 1 |
| Zt92-Zt116 | 0.049769503 | -1.5520947 | 1.6516337 | 1 |
| Zt120-Zt118 | 1.14734929 | -0.4545149 | 2.7492135 | 0.3717485 |
| Zt36-Zt118 | 0.567034576 | -1.0348296 | 2.1688988 | 0.9820156 |
| Zt55-Zt118 | 0.652260637 | -0.9496036 | 2.2541248 | 0.9513652 |
| Zt74-Zt118 | 0.241165781 | -1.3606984 | 1.84303 | 0.9999931 |
| Zt80-Zt118 | 0.571196807 | -1.0306674 | 2.173061 | 0.9810012 |
| Zt92-Zt118 | 0.589800532 | -1.0120637 | 2.1916647 | 0.9759244 |
| Zt36-Zt120 | -0.580314714 | -2.1821789 | 1.0215495 | 0.9786267 |
| Zt55-Zt120 | -0.495088653 | -2.0969529 | 1.1067756 | 0.9938306 |
| Zt74-Zt120 | -0.906183509 | -2.5080477 | 0.6956807 | 0.7063957 |
| Zt80-Zt120 | -0.576152483 | -2.1780167 | 1.0257117 | 0.9797371 |
| Zt92-Zt120 | -0.557548758 | -2.159413 | 1.0443154 | 0.9841718 |
| Zt55-Zt36 | 0.085226062 | -1.5166381 | 1.6870903 | 1 |
| Zt74-Zt36 | -0.325868795 | -1.927733 | 1.2759954 | 0.9998592 |
| Zt80-Zt36 | 0.004162232 | -1.597702 | 1.6060264 | 1 |
| Zt92-Zt36 | 0.022765957 | -1.5790983 | 1.6246302 | 1 |
| Zt74-Zt55 | -0.411094857 | -2.0129591 | 1.1907694 | 0.9987597 |
| Zt80-Zt55 | -0.08106383 | -1.682928 | 1.5208004 | 1 |
| Zt92-Zt55 | -0.062460105 | -1.6643243 | 1.5394041 | 1 |
| Zt80-Zt74 | 0.330031027 | -1.2718332 | 1.9318952 | 0.9998408 |
| Zt92-Zt74 | 0.348634752 | -1.2532295 | 1.950499 | 0.9997307 |
| Zt92-Zt80 | 0.018603725 | -1.5832605 | 1.6204679 | 1 |

Table S10. One-way Analysis of Variance of the radius of zones of inhibition produced by *Pseudomonas* isolate Roth107 in confrontation with genetically diverse *Z. tritici* isolates in the blastospore antagonism assay *in vitro*.

|  | Df | Sum Sq | Mean Sq | *F* value | Pr(>*F*) |  |
| --- | --- | --- | --- | --- | --- | --- |
| *Z. tritici* genotype | 11 | 23.32 | 2.1203 | 5.567 | 3.94E-05 | *** |
| Residuals | 36 | 13.71 | 0.3809 |  |  |  |

Significance codes: 0 ‘***’ 0.001 ‘**’ 0.01 ‘*’ 0.05 ‘.’ 0.1 ‘ ’ 1

Table S11. Tukey’s test for multiple comparisons at the 5% level of significance, comparing zones of inhibition produced by *Pseudomonas* isolate Roth107 in confrontation with different *Z. tritici* genotypes *in vitro*.

| Comparison | Mean Difference | Lower Bound | Upper Bound | Padj |
| --- | --- | --- | --- | --- |
| L951-IPO323 | -0.27970745 | -1.802888542 | 1.24347365 | 0.9999484 |
| Zt10-IPO323 | -0.76655142 | -2.289732514 | 0.75662968 | 0.8297251 |
| Zt114-IPO323 | 1.24400709 | -0.279174005 | 2.76718819 | 0.2020985 |
| Zt116-IPO323 | -1.09820035 | -2.62138145 | 0.42498074 | 0.3623116 |
| Zt118-IPO323 | 0.54451241 | -0.978668684 | 2.06769351 | 0.9806394 |
| Zt120-IPO323 | 0.25153369 | -1.271647407 | 1.77471478 | 0.9999822 |
| Zt36-IPO323 | -1.18480496 | -2.70798606 | 0.33837613 | 0.2597812 |
| Zt55-IPO323 | 0.69613918 | -0.827041911 | 2.21932028 | 0.8997142 |
| Zt74-IPO323 | 0.09932181 | -1.423859286 | 1.62250291 | 1 |
| Zt80-IPO323 | -0.25952571 | -1.782706805 | 1.26365539 | 0.9999756 |
| Zt92-IPO323 | 0.31906472 | -1.204116379 | 1.84224581 | 0.9998132 |
| Zt10-L951 | -0.48684397 | -2.010025068 | 1.03633712 | 0.99189 |
| Zt114-L951 | 1.52371454 | 0.000533441 | 3.04689563 | 0.0498527 |
| Zt116-L951 | -0.81849291 | -2.341674005 | 0.70468819 | 0.7657594 |
| Zt118-L951 | 0.82421986 | -0.698961238 | 2.34740095 | 0.7581682 |
| Zt120-L951 | 0.53124113 | -0.991939962 | 2.05442223 | 0.983925 |
| Zt36-L951 | -0.90509752 | -2.428278614 | 0.61808358 | 0.6428864 |
| Zt55-L951 | 0.97584663 | -0.547334466 | 2.49902773 | 0.5362282 |
| Zt74-L951 | 0.37902926 | -1.144151841 | 1.90221035 | 0.9990618 |
| Zt80-L951 | 0.02018174 | -1.50299936 | 1.54336283 | 1 |
| Zt92-L951 | 0.59877216 | -0.924408933 | 2.12195326 | 0.9616884 |
| Zt114-Zt10 | 2.01055851 | 0.487377413 | 3.53373961 | 0.0025229 |
| Zt116-Zt10 | -0.33164894 | -1.854830033 | 1.19153216 | 0.9997296 |
| Zt118-Zt10 | 1.31106383 | -0.212117266 | 2.83424493 | 0.1488991 |
| Zt120-Zt10 | 1.01808511 | -0.50509599 | 2.5412662 | 0.4734858 |
| Zt36-Zt10 | -0.41825355 | -1.941434642 | 1.10492755 | 0.9977402 |
| Zt55-Zt10 | 1.4626906 | -0.060490494 | 2.9858717 | 0.0694378 |
| Zt74-Zt10 | 0.86587323 | -0.657307869 | 2.38905432 | 0.7003882 |
| Zt80-Zt10 | 0.50702571 | -1.016155388 | 2.0302068 | 0.9887936 |
| Zt92-Zt10 | 1.08561613 | -0.437564962 | 2.60879723 | 0.3788678 |
| Zt116-Zt114 | -2.34220745 | -3.865388542 | -0.81902635 | 0.0002711 |
| Zt118-Zt114 | -0.69949468 | -2.222675775 | 0.82368642 | 0.8968586 |
| Zt120-Zt114 | -0.9924734 | -2.515654499 | 0.53070769 | 0.511317 |
| Zt36-Zt114 | -2.42881206 | -3.951993151 | -0.90563096 | 0.0001496 |
| Zt55-Zt114 | -0.54786791 | -2.071049003 | 0.97531319 | 0.9797327 |
| Zt74-Zt114 | -1.14468528 | -2.667866378 | 0.37849581 | 0.304686 |
| Zt80-Zt114 | -1.5035328 | -3.026713897 | 0.0196483 | 0.0557038 |
| Zt92-Zt114 | -0.92494237 | -2.448123471 | 0.59823872 | 0.6131166 |
| Zt118-Zt116 | 1.64271277 | 0.119531671 | 3.16589386 | 0.0252676 |
| Zt120-Zt116 | 1.34973404 | -0.173447053 | 2.87291514 | 0.1236805 |
| Zt36-Zt116 | -0.08660461 | -1.609785705 | 1.43657649 | 1 |
| Zt55-Zt116 | 1.79433954 | 0.271158443 | 3.31752064 | 0.0100835 |
| Zt74-Zt116 | 1.19752216 | -0.325658932 | 2.72070326 | 0.2465233 |
| Zt80-Zt116 | 0.83867465 | -0.684506451 | 2.36185574 | 0.7385999 |
| Zt92-Zt116 | 1.41726507 | -0.105916025 | 2.94044617 | 0.0881062 |
| Zt120-Zt118 | -0.29297872 | -1.81615982 | 1.23020237 | 0.9999185 |
| Zt36-Zt118 | -1.72931738 | -3.252498472 | -0.20613628 | 0.0150473 |
| Zt55-Zt118 | 0.15162677 | -1.371554324 | 1.67480787 | 0.9999999 |
| Zt74-Zt118 | -0.4451906 | -1.968371699 | 1.07799049 | 0.9961317 |
| Zt80-Zt118 | -0.80403812 | -2.327219218 | 0.71914297 | 0.7844774 |
| Zt92-Zt118 | -0.2254477 | -1.748628791 | 1.2977334 | 0.9999942 |
| Zt36-Zt120 | -1.43633865 | -2.959519748 | 0.08684244 | 0.0797977 |
| Zt55-Zt120 | 0.4446055 | -1.0785756 | 1.96778659 | 0.9961746 |
| Zt74-Zt120 | -0.15221188 | -1.675392975 | 1.37096922 | 0.9999999 |
| Zt80-Zt120 | -0.5110594 | -2.034240494 | 1.0121217 | 0.9880756 |
| Zt92-Zt120 | 0.06753103 | -1.455650068 | 1.59071212 | 1 |
| Zt55-Zt36 | 1.88094415 | 0.357763052 | 3.40412524 | 0.0058415 |
| Zt74-Zt36 | 1.28412677 | -0.239054323 | 2.80730787 | 0.1687723 |
| Zt80-Zt36 | 0.92527925 | -0.597901842 | 2.44846035 | 0.6126089 |
| Zt92-Zt36 | 1.50386968 | -0.019311416 | 3.02705078 | 0.0556013 |
| Zt74-Zt55 | -0.59681737 | -2.119998471 | 0.92636372 | 0.9625435 |
| Zt80-Zt55 | -0.95566489 | -2.47884599 | 0.5675162 | 0.5666825 |
| Zt92-Zt55 | -0.37707447 | -1.900255564 | 1.14610663 | 0.999105 |
| Zt80-Zt74 | -0.35884752 | -1.882028615 | 1.16433358 | 0.9994327 |
| Zt92-Zt74 | 0.21974291 | -1.303438189 | 1.742924 | 0.9999955 |
| Zt92-Zt80 | 0.57859043 | -0.94459067 | 2.10177152 | 0.9698637 |

Table S12. One-way Analysis of Variance of the radius of zones of inhibition produced by *Pseudomonas* isolate Roth112 in confrontation with genetically diverse *Z. tritici* isolates in the blastospore antagonism assay *in vitro*.

|  | Df | Sum Sq | Mean Sq | *F* value | Pr(>*F*) |  |
| --- | --- | --- | --- | --- | --- | --- |
| *Z. tritici* genotype | 11 | 63.63 | 5.785 | 8.11 | 6.92E-07 | *** |
| Residuals | 36 | 25.68 | 0.713 |  |  |  |

Significance codes: 0 ‘***’ 0.001 ‘**’ 0.01 ‘*’ 0.05 ‘.’ 0.1 ‘ ’ 1

Table S13. Tukey’s test for multiple comparisons at the 5% level of significance, comparing zones of inhibition produced by *Pseudomonas* isolate Roth112 in confrontation with different *Z. tritici* genotypes *in vitro*.

| Comparison | Mean Difference | Lower Bound | Upper Bound | Padj |
| --- | --- | --- | --- | --- |
| L951-IPO323 | 0.663989363 | -1.4203954 | 2.7483741 | 0.9921064 |
| Zt10-IPO323 | -1.180443262 | -3.264828 | 0.9039415 | 0.7050454 |
| Zt114-IPO323 | 1.284955675 | -0.7994291 | 3.3693404 | 0.5919036 |
| Zt116-IPO323 | -1.653439715 | -3.7378245 | 0.4309451 | 0.2356942 |
| Zt118-IPO323 | 1.950270389 | -0.1341144 | 4.0346552 | 0.0845742 |
| Zt120-IPO323 | 0.671023937 | -1.4133608 | 2.7554087 | 0.9914071 |
| Zt36-IPO323 | -1.308847517 | -3.3932323 | 0.7755373 | 0.5654938 |
| Zt55-IPO323 | -1.004388298 | -3.0887731 | 1.0799965 | 0.8647011 |
| Zt74-IPO323 | -1.6191578 | -3.7035426 | 0.465227 | 0.2614746 |
| Zt80-IPO323 | -1.004104609 | -3.0884894 | 1.0802802 | 0.8649097 |
| Zt92-IPO323 | -0.415172872 | -2.4995576 | 1.6692119 | 0.9998853 |
| Zt10-L951 | -1.844432625 | -3.9288174 | 0.2399521 | 0.1248323 |
| Zt114-L951 | 0.620966312 | -1.4634185 | 2.7053511 | 0.9954551 |
| Zt116-L951 | -2.317429078 | -4.4018138 | -0.2330443 | 0.0186877 |
| Zt118-L951 | 1.286281026 | -0.7981037 | 3.3706658 | 0.5904391 |
| Zt120-L951 | 0.007034574 | -2.0773502 | 2.0914193 | 1 |
| Zt36-L951 | -1.97283688 | -4.0572217 | 0.1115479 | 0.0776012 |
| Zt55-L951 | -1.668377661 | -3.7527624 | 0.4160071 | 0.2250374 |
| Zt74-L951 | -2.283147163 | -4.3675319 | -0.1987624 | 0.0217046 |
| Zt80-L951 | -1.668093972 | -3.7524787 | 0.4162908 | 0.2252366 |
| Zt92-L951 | -1.079162235 | -3.163547 | 1.0052225 | 0.8037089 |
| Zt114-Zt10 | 2.465398937 | 0.3810142 | 4.5497837 | 0.0096369 |
| Zt116-Zt10 | -0.472996453 | -2.5573812 | 1.6113883 | 0.9996003 |
| Zt118-Zt10 | 3.130713651 | 1.0463289 | 5.2150984 | 0.0003931 |
| Zt120-Zt10 | 1.851467199 | -0.2329176 | 3.935852 | 0.1217361 |
| Zt36-Zt10 | -0.128404255 | -2.212789 | 1.9559805 | 1 |
| Zt55-Zt10 | 0.176054964 | -1.9083298 | 2.2604397 | 1 |
| Zt74-Zt10 | -0.438714539 | -2.5230993 | 1.6456702 | 0.9998044 |
| Zt80-Zt10 | 0.176338653 | -1.9080461 | 2.2607234 | 1 |
| Zt92-Zt10 | 0.76527039 | -1.3191144 | 2.8496552 | 0.9764222 |
| Zt116-Zt114 | -2.93839539 | -5.0227802 | -0.8540106 | 0.0010174 |
| Zt118-Zt114 | 0.665314714 | -1.4190701 | 2.7496995 | 0.9919783 |
| Zt120-Zt114 | -0.613931738 | -2.6983165 | 1.470453 | 0.9958709 |
| Zt36-Zt114 | -2.593803192 | -4.678188 | -0.5094184 | 0.0053245 |
| Zt55-Zt114 | -2.289343972 | -4.3737287 | -0.2049592 | 0.0211277 |
| Zt74-Zt114 | -2.904113475 | -4.9884982 | -0.8197287 | 0.0012034 |
| Zt80-Zt114 | -2.289060284 | -4.3734451 | -0.2046755 | 0.0211538 |
| Zt92-Zt114 | -1.700128547 | -3.7845133 | 0.3842562 | 0.2035446 |
| Zt118-Zt116 | 3.603710104 | 1.5193253 | 5.6880949 | 0.0000364 |
| Zt120-Zt116 | 2.324463652 | 0.2400789 | 4.4088484 | 0.0181192 |
| Zt36-Zt116 | 0.344592198 | -1.7397926 | 2.428977 | 0.999982 |
| Zt55-Zt116 | 0.649051418 | -1.4353334 | 2.7334362 | 0.9934405 |
| Zt74-Zt116 | 0.034281915 | -2.0501029 | 2.1186667 | 1 |
| Zt80-Zt116 | 0.649335106 | -1.4350497 | 2.7337199 | 0.993417 |
| Zt92-Zt116 | 1.238266843 | -0.8461179 | 3.3226516 | 0.6432205 |
| Zt120-Zt118 | -1.279246452 | -3.3636312 | 0.8051383 | 0.59821 |
| Zt36-Zt118 | -3.259117906 | -5.3435027 | -1.1747331 | 0.0002069 |
| Zt55-Zt118 | -2.954658687 | -5.0390435 | -0.8702739 | 0.0009394 |
| Zt74-Zt118 | -3.569428189 | -5.653813 | -1.4850434 | 0.0000433 |
| Zt80-Zt118 | -2.954374998 | -5.0387598 | -0.8699902 | 0.0009407 |
| Zt92-Zt118 | -2.365443261 | -4.449828 | -0.2810585 | 0.0151161 |
| Zt36-Zt120 | -1.979871454 | -4.0642562 | 0.1045133 | 0.0755318 |
| Zt55-Zt120 | -1.675412234 | -3.759797 | 0.4089725 | 0.22014 |
| Zt74-Zt120 | -2.290181737 | -4.3745665 | -0.205797 | 0.0210509 |
| Zt80-Zt120 | -1.675128546 | -3.7595133 | 0.4092562 | 0.220336 |
| Zt92-Zt120 | -1.086196809 | -3.1705816 | 0.998188 | 0.7973926 |
| Zt55-Zt36 | 0.30445922 | -1.7799256 | 2.388844 | 0.9999949 |
| Zt74-Zt36 | -0.310310283 | -2.3946951 | 1.7740745 | 0.9999938 |
| Zt80-Zt36 | 0.304742908 | -1.7796419 | 2.3891277 | 0.9999949 |
| Zt92-Zt36 | 0.893674645 | -1.1907101 | 2.9780594 | 0.9320226 |
| Zt74-Zt55 | -0.614769503 | -2.6991543 | 1.4696153 | 0.9958231 |
| Zt80-Zt55 | 0.000283689 | -2.0841011 | 2.0846685 | 1 |
| Zt92-Zt55 | 0.589215426 | -1.4951693 | 2.6736002 | 0.9970924 |
| Zt80-Zt74 | 0.615053192 | -1.4693316 | 2.699438 | 0.9958068 |
| Zt92-Zt74 | 1.203984928 | -0.8803998 | 3.2883697 | 0.6802073 |
| Zt92-Zt80 | 0.588931737 | -1.495453 | 2.6733165 | 0.9971044 |
